# Immune-based mutation classification enables neoantigen prioritization and immune feature discovery in cancer immunotherapy

**DOI:** 10.1101/700732

**Authors:** Peng Bai, Yongzheng Li, Qiuping Zhou, Jiaqi Xia, Peng-Cheng Wei, Hexiang Deng, Min Wu, Sanny K. Chan, John W. Kappler, Yu Zhou, Eric Tran, Philippa Marrack, Lei Yin

## Abstract

Genetic mutations lead to the production of mutated proteins from which peptides are presented to T cells as cancer neoantigens. Evidence suggests that T cells that target neoantigens are the main mediators of effective cancer immunotherapies. Although algorithms have been used to predict neoantigens, only a minority are immunogenic. The factors that influence neoantigen immunogenicity are not completely understood. Here, we classified human neoantigen/neopeptide data into three categories based on their TCR-pMHC binding events. We observed a conservative mutant orientation of the anchor residue from immunogenic neoantigens which we termed the “NP” rule. By integrating this rule with an existing prediction algorithm, we found improved performance in neoantigen prioritization. To better understand this rule, we solved several neoantigen/MHC structures. These structures showed that neoantigens that follow this rule not only increase peptide-MHC binding affinity but also create new TCR-binding features. These molecular insights highlight the value of immune-based classification in neoantigen studies and may enable the design of more effective cancer immunotherapies.

## Introduction

The immune system can recognize and destroy cancer cells^1^ and CD8^+^ T cells are a major component in this process ^2–5^. The recognition of tumor cells by CD8+ T cells requires at least two components: expression of major histocompatibility complex class I proteins (MHCI) on the cancer cells and tumor-derived antigenic peptides which bind to MHCs. T-cell receptors (TCRs) expressed by T cells recognize these peptide-MHC complexes (pMHC) on cancer cells and target them for destruction.

During the last 25 years, great effort has been made to identify the tumor antigens that are targeted by T cells^6^. Cancer antigens can be classified into two broad categories, self-antigens which are expressed by some normal tissues but are usually expressed at much higher levels by cancer cells (e.g., MART-1, HER2, CEA) and nonself-antigens, which include antigens derived from oncogenic viruses (e.g., human papilloma virus), or neoantigens derived from somatic mutations in cancer cells^7^. Neoantigens are not present in normal tissues, hence the immune system is not tolerant to them and views them as foreign antigens ^8,9^. The targeting of neoantigens is ideal because they are not expressed by normal tissue and therefore should not result in “on-target, off-tumor” toxicity which can be observed in immunotherapies that target self antigens. Moreover, unlike self antigens, neoantigens are foreign to the immune system and thus highly restricted TCRs should exist in patients (i.e., not deleted during thymic selection).

Currently, the identification of neoantigen relies on next-generation sequencing (NGS) to identify nonsynonymous mutations, followed by predicting the theoretical binding of the corresponding mutated peptides to the patient’s HLA molecule ^10–16^using peptide/MHCI binding prediction algorithms. Binding predictions have been used to narrow down and select neoantigen candidates for use in immunotherapy approaches as well as to help identify the minimal epitope recognized by T cells ^7,11,12,14,17–19,19–22,22–25^. However, among the neopeptides with predicted MHC binding, only a small portion of them are immunogenic and therapeutically relevant ^26,27^. Other parameters beyond peptide-MHC binding affinity could affect neoantigen immunogenicity.

It has been hypothesized that TCR:pMHC ternary binding events may influence neoantigen immunogenicity. Yadav *et al*. and Fritsch *et al*. suggested that the side chains of neoantigen mutations pointing toward the TCR would be more immunogenic while Duan *et al*. suggested that neoantigen substitutions at MHC anchor positions may be more important^7,18,28^. Recently, Capietto *et al*. also suggested that the increased neoantigen-MHC affinity relative to the corresponding wild-type peptide is predictive of immunogenicity^29^. However, studies in this area are still limited due to the lack of large human neoantigen datasets and neoantigen-MHC crystal structures. Thus, systematic analyses based on human neoantigen data and a structural understanding of neoantigen immune properties are still needed.

Here, we attempted to determine immune features based on validated neoantigens after mutation positional classification. We found that almost all immunogenic anchor mutated neoantigens in our datasets followed a conservative mutation orientation (termed the NP rule), rather than other orientations. Combining this rule with the binding predictor NetMHCpan 4.0^24^ could improve the performance of neoantigen prioritization. To provide structural insights of this rule, we solved several pMHC structures: the KRAS G12D neoantigens in complex with HLA-C*08:02 and the ineffective mouse DPAGT1 peptides in complex with a mouse MHC (H-2 Kb). We showed that neoantigens that follow the NP rule could generate immune features for T cell recognition. Our data also suggests that the antigen exposed surface area may be associated with neoantigen immunogenicity

## Materials and Methods

### Generation of the InEffective Neopeptides Dataset (IEND)

To generate a length-fitted ineffective neopeptide dataset, mutant peptides from RIEND (Raw InEffective Neopeptides Dataset) were cleaved into 9mer and 10mer containing mutations *in silico* and performed binding prediction with the NetMHCpan 4.0 server^24^. This newly generated dataset was called the InEffective Neopeptide Dataset (IEND).

The mutant peptides (9mer and 10mer) were inputted in the NetMHCpan 4.0 Server with custom python scripts. The prediction of immunogenic neoantigens relies on a recommend cut-off (IC50 < 500 nM) of a predicted MHCI binding affinity.

### Comparison of amino acid biochemical properties

Amino acid dissimilarity comparison of wild-type and mutant amino acids was taken for peptide pairs between immunogenic and ineffective cohorts based on the BLOSUM50 matrix^30^. This matrix provided BLOSUM scores represented the similarity of amino acid pairs (higher score indicates more similar). Tails of the violins to the range of the data were not trimmed. The figure was generated by R. Statistical analyses were calculated by Wilcoxon test.

The comparisons of”hydrophobicity”, “polarity”, and “side chain bulkiness” scores of mutant amino acids between immunogenic and ineffective cohorts were taken based on independent numeric scales^31^. Independent numeric scales of each property were recorded in Supplementary Table 1, 3, and 4. Related figures were generated by R. Statistical analyses for the “hydrophobicity”, “polarity”, and “side chain bulkiness” differences were performed using Wilcoxon test.

### Differential agretopic index (DAI) score calculation

The calculation of DAI was described previously^32^. Briefly, peptide binding affinity with HLAs was predicted by NetMHCpan 4.0. The DAI score of each neoantigen pair was calculated by subtraction of the neoantigen predicted IC50 binding affinity from the corresponding wild-type counterparts.

### Anchor, MHC-contacting, and TCR-contacting positions determination

Anchors were defined for each allele based on the SYFPEITHI database^33^ and the highest information content in the NetMHCpan binding motif record^24^. The anchor position for each entry was cross-validated based on solved HLA structures from Protein Data Bank (PDB). We recorded these results as “consensus anchor” in IND and IEND (Supplementary Table 1 and 3).

Peptide MHC-contacting and TCR-contacting positions were determined based on solved peptide-MHC complex structures. Briefly, the positions of peptides that were proved to be non-anchor positions can be divided into MHC-contacting and TCR-contacting positions. Based on pMHC structural models from PDB, those positions which contact MHC were treated as MHC-contacting positions. On the contrary, TCR-contacting positions often harbor residues with side chains that point toward outside from the pMHC complex and may contact with TCRs. The MHC-contacting and TCR-contacting information of different HLAs was recorded in IND and IEND.

### Antigen library and the determination of HLA anchor position preference

The nonameric peptide libraries of 30 HLA alleles were obtained from the IEDB database^34^. Sequence logos were generated using the sequence logo generator^35^. The threshold for preferential amino acids at anchor positions was set to include and above 10% based on the nonameric peptide libraries from IEDB. Related information was recorded in Supplementary Table 1 and 3.

### Combined NP+Binding (Con NP+B) model building

To combine the NP rule with NetMHCpan 4.0, we used a logistic regression algorithm. Analyses were performed using the R build-in function glm(), as below:

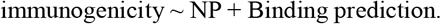

Anchor mutated neoantigen data were selected to train the model. The performance of this model was shown by the ROC curve.

The ROC curve was plotted from the false positive rate (FPR) and true positive rate (TPR) values calculated by varying the cut-off value (separating the predicted positive from the predicted negative) from high to low. The plots were generated in R using the packages ggplot2 and plotROC.

To further test this model, we resampled 50 times. Random resampling of the data (2/3rd resampling) was used for training. The AUC values were calculated by plotROC package. After iteration, the differences of AUC between the four models were measured using paired t test in R.

### Protein Expression, refolding, and purification

Inclusion bodies of HLA heavy chains and β2M were expressed as described previously^36^. Briefly, The DNA encoding MHC heavy chain (HLA-C*08:02 and H-2 Kb) and light chain (human β2M and mouse β2M) was synthesized (Idobio) and cloned into pET-22(b) vector (Novagen). The vectors were transformed into the E. coli strain BL21 DE3 (Novagen). Transformants were selected on Luria broth (LB) agar plates containing ampicillin. A single colony was selected and cultured in LB fluid medium with the antibiotics listed above at 37°C. Upon reaching an optical density OD600 of 0.6, expression was induced with the addition of 1mM IPTG. Incubation continued at 37°C for 5h. The cells were harvested by centrifugation and then resuspended in PBS buffer with 1 mM PMSF at 4°C. The cells were lysed, and the lysate was clarified by centrifugation at 10,000 g to collect inclusion bodies. Inclusion bodies were harvested and solubilized in 20 mM Tris (Vetec) pH 8.0, 8 M urea (Vetec), 1 mM EDTA (BBI life sciences), 1 mM DTT (Sinopharm chemical reagent) and 0.2 mM PMSF (Sinopharm chemical reagent).

Refolding was performed in the presence of MHC heavy chain, β2M, and peptides as described previously^37^. Briefly, the resolubilized heavy chain (60 mg each) and the light chain (25 mg each) in the presence of the corresponding peptide were added into 1 liter of refolding buffer [100 mM Tris (pH 8.4), 0.5 mM oxidized glutathione (BBI life sciences), 5 mM reduced glutathione (BBI life sciences), 400mM L-arginine (Vetec), 2mM EDTA (BBI life sciences)]. After 48 h of refolding, the 1 L mixture was transferred into dialysis bags (Spectra) and dialyzed against 15 liters of 10 mM Tris buffer (pH 8.0) at 4°C for 24h.

Refolded proteins were purified by anion exchange chromatography with Q Sepharose HP (GE Healthcare) column then Mono Q column (GE Healthcare) and concentrated by tangential flow filtration using Amicon Ultra centrifugal filters (Merck). For desalination and purification, samples were loaded onto a Superdex 200 increase 10/300 GL column (GE Healthcare) for size exclusion chromatography. Chromatography was taken with BioLogic DuoFlow system (Bio-rad) at a flow rate of 1 mL/min. Peak analysis was performed using the ASTRA software package (BioLogic Chromatography Systems).

### Thermal stability assay

The thermal stability assay was performed in the Real Time Detection system (Roche). Each pMHC complex was diluted in 10 mM Tris-HCl (pH 8.0), 0.1 M NaCl buffer. The experiment was performed in triplicates for different pMHC complexes. Both pMHC complexes were heated from 10 to 95 °C with a heating rate of 1 °C per minute. The fluorescence intensity was measured with excitation at 530 nm and emission at 555 nm. The Tm represents the temperature for which 50% of the protein is unfolded.

### Crystallization, data collection, and processing

Purified pMHC complexes were concentrated to 10 mg/ml for crystallization trials before screening using a series of kits from Hampton Research. Protein complexes were crystallized by sitting drop vapor diffusion technique at 4 °C. Single crystals of C08-mut9m and C08-mut10m were obtained in the condition of 0.2 M ammonium acetate, 0.1 M HEPES (pH 6.5), 25% w/v polyethylene glycol 3,350. For the H-2 Kb complex, single crystals of Kb-8mV and Kb-8mL complex were obtained when 4% v/v Tacsimate (pH 6.0), 12% w/v Polyethylene glycol 3,350 was used as the reservoir buffer.

Crystals were transferred to crystallization buffer containing 20% (w/v) glycerol and flash-cooled in liquid nitrogen immediately. The diffraction data were collected at the Shanghai Synchrotron Radiation Facility (Shanghai, China) on beamline BL17U1/BL18U1/BL19U1, and processed using the iMosflm program^38^. Data reduction was performed with Aimless and Pointless in the CCP4 software suite^39^. All structures were determined by molecular replacement using Phaser^40^. The models from the molecular replacement were built using the COOT (Crystallographic Object-Oriented Toolkit) program^41^ and subsequently subjected to refinement using Phenix software^42^. Data collection, processing, and refinement statistics are summarized in Supplementary Table 7. All the structural figures were prepared using PyMOL (http://www.pymol.org) program. The atomic coordinates and structure factors for the reported crystal structures have been deposited on the Protein Data Bank (PDB; http://www.rcsb.org/pdb/).

## Results

### Classification of cancer neoantigen/neopeptide data by mutation position

We obtained MHCI-related, immunogenic (positive data, termed neoantigen) and ineffective (negative data, termed neopeptide) clinical human cancer neoantigen/neopeptide data from a public database called NEPdb (http://nep.whu.edu.cn). Herein, the mutant peptides which proved to induce T cell response or led to clinical response in the context of certain HLA are termed ‘neoantigen’. The mutant peptides which did not induce T cell or clinical response are termed ‘neopeptide’. The neoantigen-HLA complex is termed ‘neoepitope’. These data were curated from 34 published studies that contain neoantigen/neopeptide sequences and relevant HLAs. All of the data were tested in *in vitro* T cell assays or clinical therapies^5,43,12–14,11,21,44–54,17,55,56,19,57–62,22,63,64,65(p53).^

The Immunogenic Neoantigen Dataset (IND) in this study, contains 128 neoantigens with corresponding MHCIs (Fig. 1a; Supplementary Table 1). The Raw InEffective Neopeptide Dataset (RIEND) contains 11739 neopeptides with corresponding MHCIs (Fig. 1a; Supplementary Table 2). All of these mutant peptides had been proven to be ineffective. The majority of them are not in the MHCI-fitted lengths (most MHCI binders are 8-11 amino acids in length). To obtain negative control for further study, we processed these raw ineffective neopeptides into optimal MHC-binding length (9mer and 10mer) *in silico* and used the NetMHCpan 4.0 algorithm for peptide-MHC binding prediction. This processed dataset is termed the InEffective Neopeptide Dataset (IEND) and contains 2883 entries (Fig. 1a; Supplementary Table 3).

**Figure 1.**
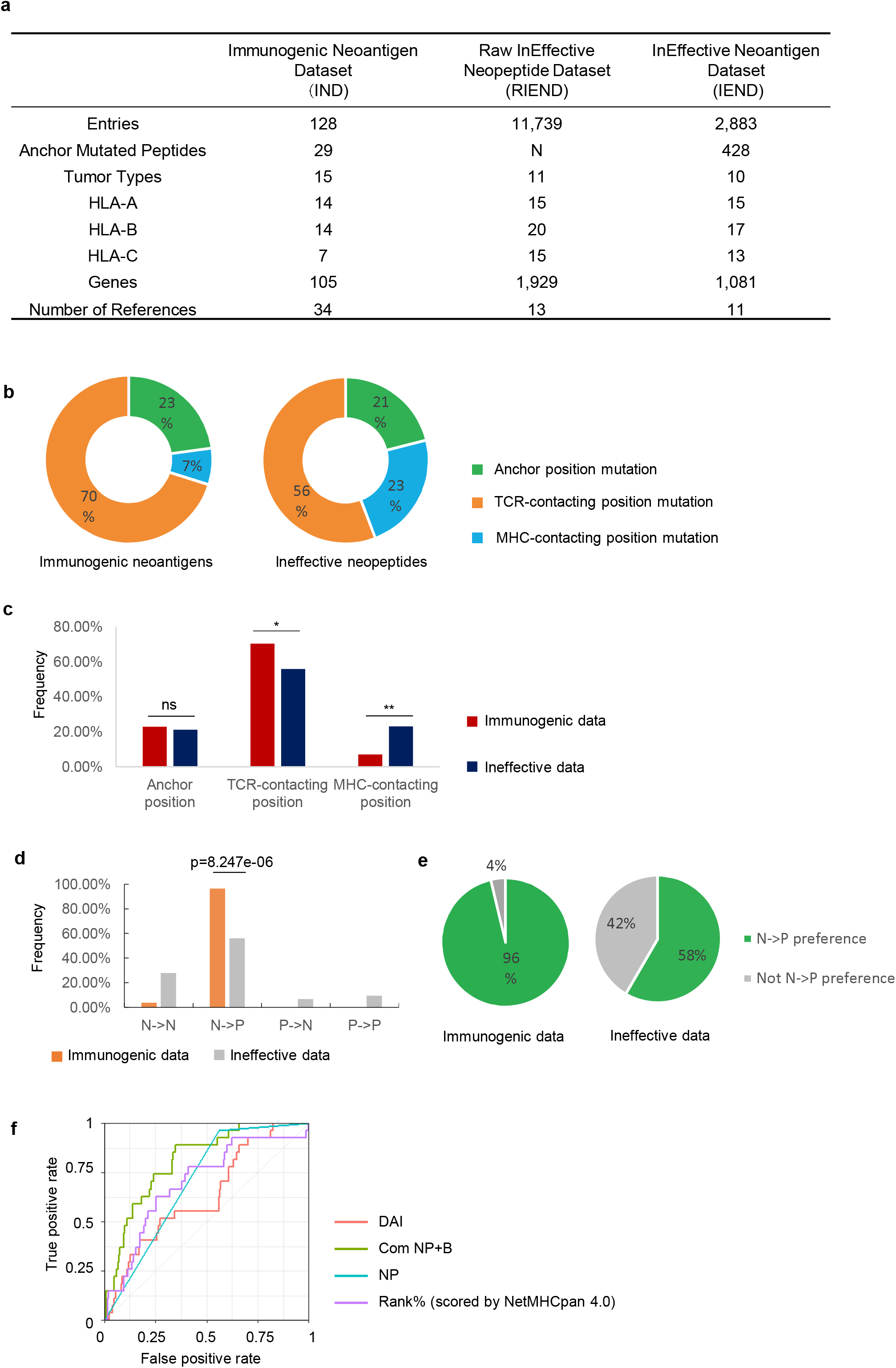
Immune-based classification of data from the Immunogenic neoantigen dataset (IND) and Ineffective neopeptides dataset (IEND). **a**, Overview of the IND, RIEND, and IEND datasets. **b**, Donut plots of the percentage of 9mer neoantigen/neopeptides, categorized into three classes based on different mutation positions: anchor mutation, MHC-contacting position, and TCR-contacting position (immunogenic data, n =51; ineffective data, n=763). **c**, Nonameric peptides mutation distribution of three classes (mutated at anchor mutation, MHC-contacting position, and TCR-contacting position) from IND and IEND. The frequency of mutation distribution at TCR-contacting position and MHC-contacting position showed significant difference (TCR-contacting position, p=0.0378; MHC-contacting position, P=0.0027. n (immunogenic) =51; n (ineffective) =763. Fisher’s exact test). The percentage of mutation distribution at the anchor position did not show a significant difference between immunogenic and ineffective data (ns, non-significant). **d**, Nonameric peptides distribution of four subgroups (NN, NP, PN, and PP). A significant difference was observed in the NP group (p=8.247e-06, n=27 (immunogenic); n=425 (ineffective), Fisher’s exact test). **e**, Pie charts represented the percentage of the NP group and non-NP groups in IND and IEND (n (immunogenic) = 27; n (ineffective) =425). **f**, Receiver operator characteristic (ROC) curve showed the performance of four prediction models (DAI score, binary NP rule, binding prediction (Rank% scored by NetMHCpan 4.0), combination of NP rule + binding prediction (Com NP+B)) with anchor mutated data (data from IND and IEND, n (immunogenic) = 27; n (ineffective) =425). The AUC (Area Under the ROC Curve) was calculated for each predictive model (AUC_DAI_= 0.632; AUC_NP rule_ =0.701; AUC_com NP-B_ =0.810; AUC _Rank%_=0.698).

A stable pMHC complex is helpful for effective TCR recognition and may induce T cell responses. Thus, neoepitope candidates can be prioritized through prediction algorithms by eliminating the peptides with weak binding affinity to MHCIs. While binding prediction algorithms have been successfully used for eliminating candidate neoantigens, the prediction of true positive neoantigens remains quite low^66^. Features beyond MHC binding affinity are therefore involved in neoantigen immunogenicity. Recently, some studies found that the TCR:pMHC ternary binding events may influence immunogenicity^67,68^. However, the underlying immunological mechanisms of how the binding events affect clinical outcome in patients with cancer remains poorly defined.

To address the above questions, we classified peptides from IND and IEND based on the mutation positions with different functions during TCR:pMHC binding. Specifically, mutant peptides from IND and IEND were categorized in the context of MHCs into three categories, with mutations: 1) at anchor positions that impact MHC binding; 2) at other MHC-contacting positions; and 3) that contact the TCR (i.e., point toward TCRs instead of MHCs). The classification was performed by referencing the SYFPEITHI database^33^, the NetMHCpan antigen-binding motif viewer^24^, and pMHC structures from the Protein Data Bank database (PDB). The classification information is shown in Supplementary Table 1 and 3.

We next calculated the percentage of neoantigens in the different categories. The immunogenic neoantigens were more likely to mutate at TCR-contacting regions and less likely to change at MHC-contacting regions as compared to those ineffective neopeptides (Fig. 1b, c), consistent with prior reports^7,28^. There was no difference in the frequency of neoantigen peptides that mutate at anchor positions in the immunogenic neoantigens compared to the ineffective neopeptides (Fig.1c). This suggests that the classification of anchor mutated peptides is not sufficient to distinguish the immunogenic neoantigens from the ineffective candidates.

Next, we sought to identify amino acid biochemical properties that define immunogenicity of neoantigens within the category of the “TCR-contacting mutations”. We evaluated four biochemical properties of amino acids, including “amino acid dissimilarity”, “hydrophobicity”, “polarity”, and “side chain bulkiness”, that discriminate between immunogenic and ineffective cohorts. First, we defined dissimilarity of peptide pairs (the phrase “peptide pairs” refers to the mutant and corresponding wild-type peptide), both for immunogenic and ineffective data, by using normalized BLOSUM50 substitution matrix^30^. Next, “hydrophobicity”, “polarity”, and “side chain bulkiness” properties were analyzed for mutations across immunogenic and ineffective cohorts using independent numeric scales described by *Chowell et al* (Supplementary Table. 4)^31^. We found no significant difference for the four properties between immunogenic and ineffective cohorts (Supplementary Fig. 1). Thus, in our datasets, we found that these four biochemical properties were not predictive for neoantigen immunogenicity.

### Anchor mutated neoantigens exhibit a conservative mutation pattern to acquire immunogenicity

We aimed to further characterize intrinsic immunological properties of anchor mutated neoantigens beyond binding affinity. MHC molecules have many allelic variants with different binding properties. Thus, the peptides recognized by different MHCs are very diverse, with allele-specific amino acid preferences. To investigate the preferential binding property of HLAs, we set the cut-off threshold above 10% as HLA preferential amino acids at anchor positions as described previously^69,70^ Specifically, we considered a call as preferential amino acids only if the amino acids at a certain position yielded at least a 10% enrichment, based on the binding-peptide libraries from IEDB^34^ (Supplementary Fig. 2, Supplementary Table 1 and 3). We classified the impact of a mutation at the MHC anchor residues into the following 4 groups: 1) non-preferential to non-preferential residues (NN); 2) non-preferential to preferential residues (NP); 3) preferential to non-preferential residues (PN); and 4) preferential to preferential residues (PP) (Fig. 1d; Supplementary Table 5). Statistical analysis showed a higher frequency of NP amino acid mutations in the immunogenic dataset compared to those in the ineffective dataset (Fig. 1d, p=8.247e-06). Interestingly, 26 in 27 (96%) of the immunogenic pairs can be classified into the NP group. In contrast, peptide pairs in the NN, PN, or PP groups were dramatically less immunogenic (Fig. 1e; Supplementary Table 5).

Next, we further analyzed the 1/27 case in the immunogenic cohort that did not follow the NP rule. This neoantigen (MYADM R30W, derived from the MYADM protein) has a mutation at the C-termini of the peptide^44^. Notably, the T-cell response was directed against both the wild-type and mutant MYADM peptides which demonstrated that the wild-type MYADM peptide can bind to the patient’s HLA and elicit an autoreactive T cell response *in vivo*. Neoantigens that elicit T-cell responses that target the corresponding wild-type peptide may be less desirable to target due to the potential for normal tissue targeting and tolerance. Overall, our observations suggest that the NP rule is a conservative feature of immunogenic anchor mutated neoantigens.

The NP rule of anchor mutated neoantigens thus can be treated as a binary variable (1= true NP, 0= false NP). To assess whether this variable can be used in neoantigen prioritization, we respectively tested the performance of the binary “NP” model, and the combination model of this binary model with the NetMHCpan 4.0 Rank% model (termed Com NP+B). Two prediction models, NetMHCpan 4.0 (using NetMHCpan 4.0 Rank% score) and the DAI models (differential agretopic index, the difference of predicted binding affinity between the mutated epitope and its unmutated counterpart) were also tested with our data as benchmarks. After calculation, the “Com NP+B” model achieved better performance compared with the other three models (Fig. 1f, AUC=0.810). When comparing the performance of these four predictors, an assessment was also done by 50-fold cross-validation (2/3rd random resampling) over the data to check whether the observed difference in average AUC differs significantly (Supplementary Fig. 3). Collectively, we proved that the NP rule can be used as a predictive feature to improve neoantigen prioritization.

### Anchor mutated neoantigens can generate new surface for T cell recognition

To understand how the NP rule forms the basis for understanding the immunogenicity of anchor mutated neoantigens, we attempted to solve structures of two neoepitopes containing the KRAS G12D mutations in complex with HLA-C*08:02 (C08) ^12^. Targeting of these neoepitopes with the adoptive transfer of tumor-infiltrating lymphocytes (TIL) was associated with an objective clinical response in a patient with metastatic colorectal cancer. The TIL specifically recognized the KRAS G12D 9mer neoantigen GA<u>D</u>GVGKSA and the 10mer neoantigen GA<u>D</u>GVGKSAL (both mutated at position 3 on the peptides from glycine to aspartic acid), and not the wild-type peptides, when presented by HLA-C*08:02. These two neoantigens can be categorized into the NP group (Supplementary Table 1).

To investigate the properties of these neoepitopes, we performed protein refolding for HLA-C*08:02 (C08) with four peptides: the wild-type KRAS 9mer GAGGVGKSA (wt9m); the wild-type KRAS 10mer peptide GAGGVGKSAL (wt10m); the mutant KRAS G12D 9mer peptide GA<u>D</u>GVGKSA (mut9m, mutation site is indicated with underline), and the mutant KRAS G12D 10mer peptide GA<u>D</u>GVGKSAL (mut10m). The C08-mut9m and C08-mut10m complexes were successfully obtained by refolding *in vitro*. However, the wt9m and wt10m peptides failed to refold with HLA-C*08:02 even with a ten-fold increase in the concentration of the peptides. To investigate binding details of C08-mut9m and C08-mut10m complexes, we next performed thermal denaturation tests for C08-mut9m and C08-mut10m respectively. Both pMHCs exhibited normal and similar melting points (C08-mut9m, 50±1.6 °C; C08-mut10m, 48±1.1 °C; Supplementary Table 6). These results indicated that mut9m and the mut10m can stabilize the pMHC complexes and exhibit similar stability after engaging with HLA-C*08:02.

To identify potential structural differences of C08-mut9m and C08-mut10m, we solved the crystal structures of HLA-C*08:02 in complex with mut9m at 2.4 Å (PDB id: 6JTP) and mut10m at 1.9 Å (PDB id: 6JTN) (Supplementary Table 7). Electron density for peptides was unambiguous (Fig. 2a, b). C08-mut9m and C08-mut10m showed conservative conformations except for the peptide-regions (Fig. 2c, d). Smaller residues such as alanine and serine at peptide P1 and P2 positions are preferably selected by HLA-C*08:02 (Fig. 2e, Supplementary Fig. 2) due to the narrow cleft formed by several aromatic residues (Tyr7, Phe33, Tyr67, Tyr99, Tyr59, Tyr171, Tyr159, and Trp167) which limited the size of P1 and P2 (Fig. 2f).

**Figure 2.**
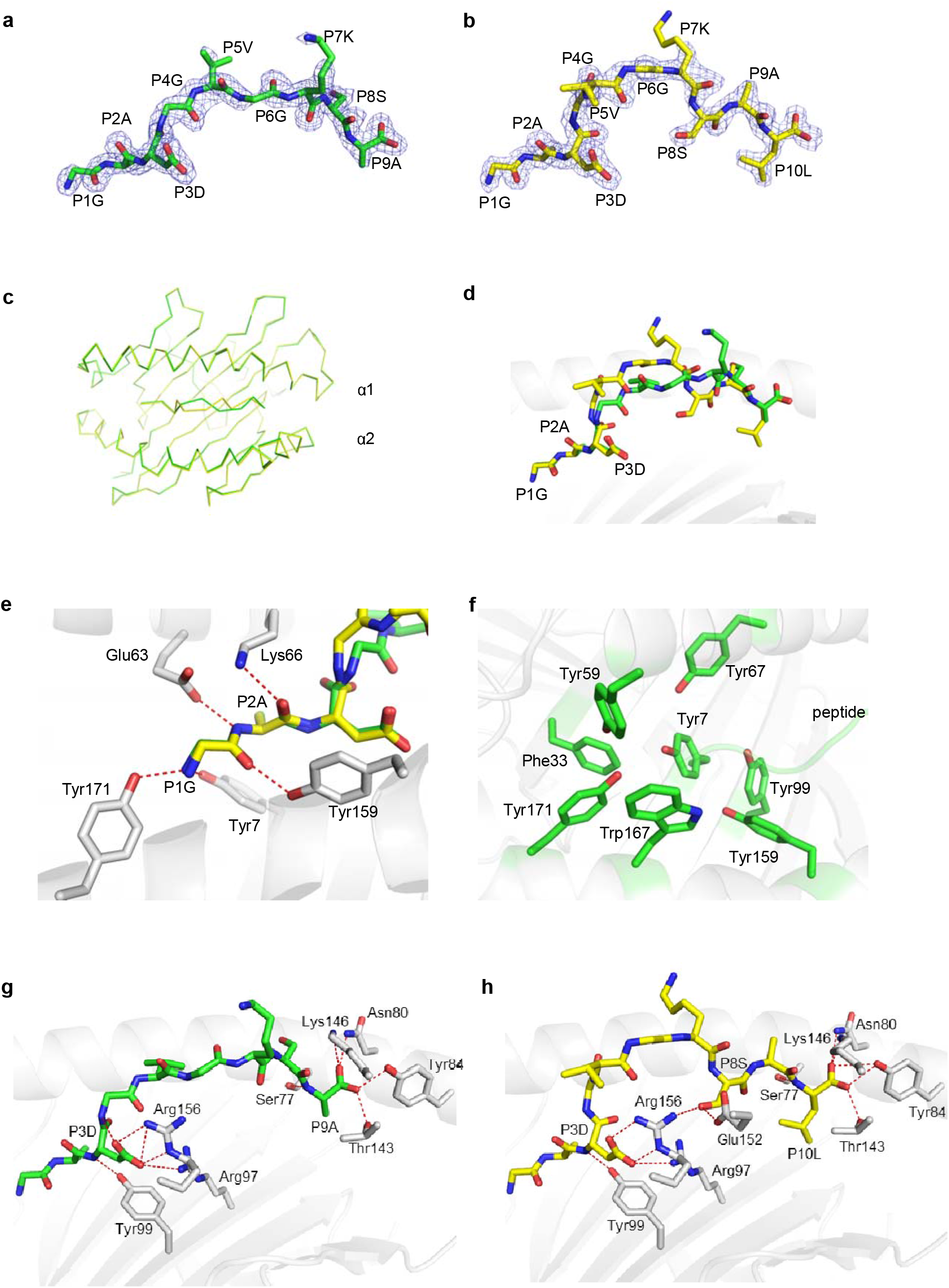
Structural comparison of the C08-mut9m and C08-mut10m complexes. **a-b**, Unambiguous 2Fo-Fc electron density maps of the (**a**) KRAS G12D 9mer (GA<u>D</u>GVGKSA, green) and (**b**)10mer (GA<u>D</u>GVGKSAL, yellow) neoantigens from the solved structures. The underlined amino acids represented the mutations. **c**, Overlay of the Cα traces (C08-mut9m, green; C08-mut10m, yellow). **d**, Overlay of the mut9m (green) and mut10m (yellow) peptides. **e**, Polar interactions at peptide P1G and P2A positions. HLA-C*08:02 showed in grey. The mut9m peptide showed in green and mut10m showed in yellow. **f**, Aromatic residues (green) from HLA-C*08:02, accommodating the P1 and P2 residues from peptides. **g**, The P3D and P9A residues of mut9m peptide interact with HLA-C*08:02. **h**, The P3D, P8S, and P10LA residues of mut10m peptide interact with HLA-C*08:02.

Generally, peptides use P2 and PΩ as anchor residues to occupy the B and F pockets of MHCIs. However, the structures of C08-mut9m and C08-mut10m revealed that mut9m and mut10m used the unconventional P3 position as an anchor to bind HLA-C*08:02 (Fig. 2g, h). HLA-C*08:02 binds P3D via the Arg97 and Arg156 residues. The side chain of P3D in C08-mut9m also forms an intra-chain hydrogen bond with P4G, while this bond is absent in C08-mut10m. Interestingly, we found that the P3D anchor residue in C08-mut9m could also provide TCR accessible surface by partially exposing its charged side chain (Supplementary Fig. 4). This phenomenon suggested that some neoantigens with anchor mutations can not only increase pMHC binding force but also provide an additional accessible surface for TCR interaction, under certain conditions, and therefore affect the interactions with TCRs.

Although mut9m and mut10m occupied the same F pocket with their PΩ residues, the P10L side chain from mut10m buried deeper than P9A in C08-mut9m (Fig. 2g, h). Meanwhile, instead of the TCR-oriented residue P8S on mut9m, mut10m uses P8S as an auxiliary anchor to bind HLAs via hydrogen bonds to Glu152 and Arg156. This residue, together with P3D, squeezed the P4-P7 region of mut10m in C08-mut10m (Fig. 2h). Together, the details of TCR-binding surfaces are largely different between C08-mut9m and C08-mut10m. We postulated that these differences may respectively modulate T cell recognition and activate different T cell repertoires (discussed below). Meanwhile, structural analysis of C08-mut9m and C08-mut10m suggested that neoantigens with anchor mutations not only generate new immune features but also create novel neoepitope surfaces. In one sense, the newly generated neoepitope can be “seen” as a totally foreign epitope by T cells. We postulated that the anchor mutated neoantigens may be more immunogenic than those neoantigens with non-anchor mutations.

### Structure of a non-therapeutic neoantigen from DPAGT1 in complex with mouse H-2 Kb

Yadav *et al* described a neopeptide that can be presented by a mouse MHC (H-2 Kb) but showed the ineffective property *in vivo*^7^. This mouse DPAGT1 V213L neopeptide contains a mutated C-terminal anchor residue that falls into the “preferential to preferential residues (PP)” group, with the changing of valine (V) to leucine (L). We next solved the X-ray crystal structures of these peptides in complex with H-2 Kb. Soluble mouse H-2 Kb in complex with the mutant DPAGT1 V213L 8mer peptide (SIIVFNL<u>L</u>, termed mut8mL) and the wild-type 8mer counterpart (SIIVFNLV, termed wt8mV) were separately expressed, refolded, and purified for crystallization trials. Crystal diffraction data of Kb-wt8mV and Kb-mut8mL were processed to 2.4 Å and 2.5 Å resolution respectively (Supplementary Table 7) and provided electron density for each peptide (Fig. 3a).

**Figure 3.**
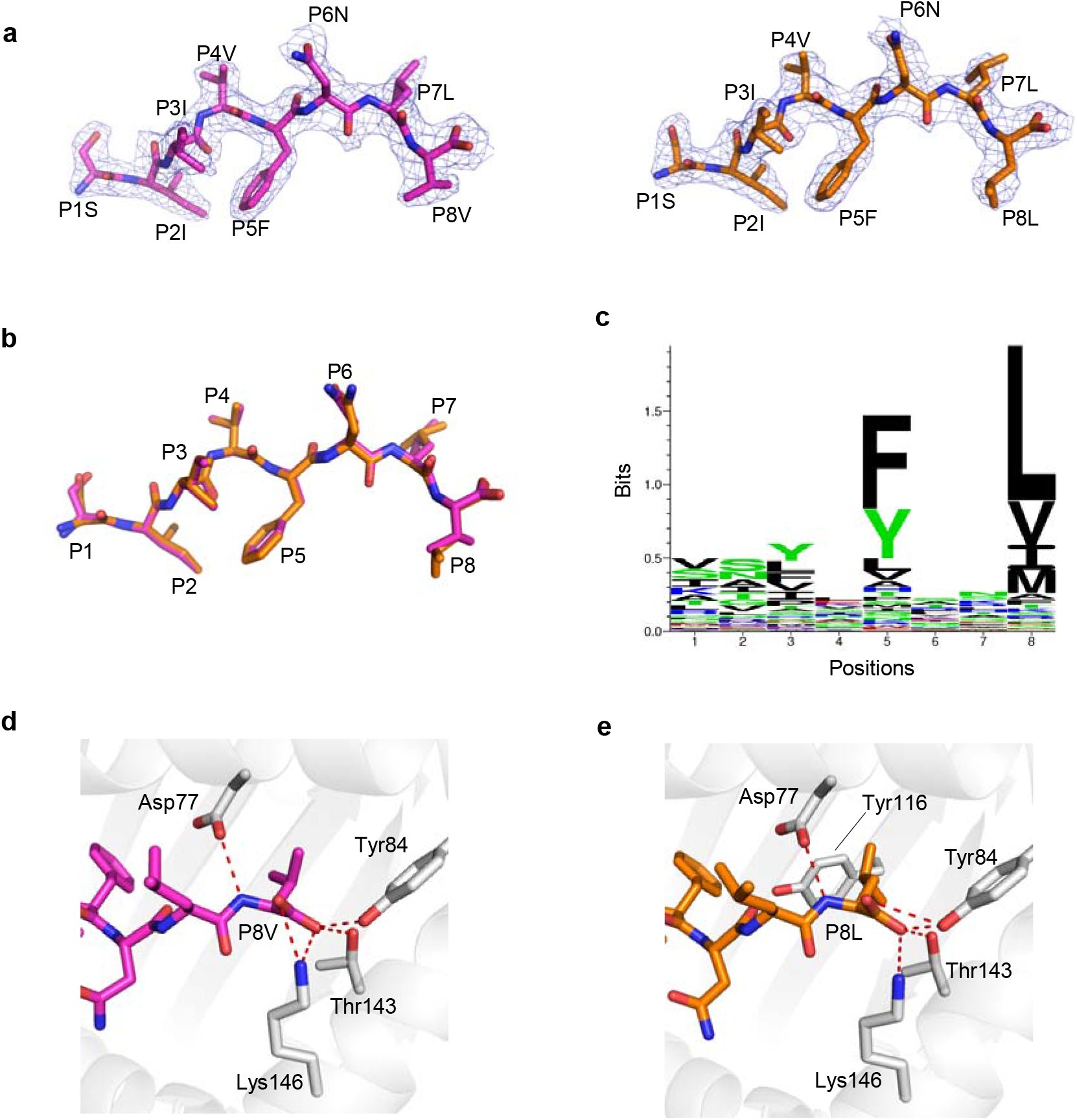
H-2 Kb presented DPAGT V213L wild type peptide wt8mV and mutant peptide mut8mL in a similar manner. **a**, Unambiguous 2Fo-Fc electron density maps of the DPAGT1 wild type 8mer peptide (wt8mV peptide SIIVFNLV, magenta) and mutant 8mer peptide (mut8mL peptide SIIVFNL<u>L</u>, orange) from solved structures. **b**, Overlay of the wt8mV (magenta) and mutant mut8mL (orange) peptides. **c**, Sequence logo of H-2 Kb with 8mer peptides. The peptide library was obtained from IEDB (n=4141). **d**, The anchor residue P8V of wt8mV interacts with H-2 Kb (grey). **e**, The anchor residue P8L of mut8mL interacts with H-2 Kb (grey).

The overall structure of Kb-wt8mV closely resembles that of Kb-mut8mL except for a slight difference at PΩ (P8) (Fig. 3b). The PΩ residues acted as anchors in both Kb-wt8mV and Kb-mut8mL (Fig. 3b). Moreover, both PΩ valine and leucine were preferably selected by H-2 Kb (Fig. 3c). Both of the two PΩ residues in these structures formed hydrogen bonds with Asp77, Tyr84, Thr143, and Lys146 (Fig. 3d, e). Although the side chain of leucine in mut8mL inserted deeper into Kb than valine in wt8mV because of its longer side chain, these two peptides did not provide a different TCR binding surface with Kb (Fig. 3b, d, e). These findings indicated that the neopeptides with the “PP” rule cannot readily change binding surfaces and be immunogenic. Ineffectiveness of the neopeptides with the PP rule might be explained by the pre-existence of the wild-type peptide-MHC complex in the thymus, which leads to negative selection of potential neopeptide restricted T cell repertoires. Considering that peptide-MHC binding is necessary for neoepitope immunogenicity, we do not discuss the situations of the “PN” and “NN” rules.

### Neoantigen exposed surface areas may affect T cell selection in cancer immunotherapy

Peptide antigens can form stable complexes with HLAs by lying within the HLA antigen-binding cleft. Some antigens, called “featureless” antigens, have a relatively small exposed surface area (ESA) from side chains pointing towards the T cell receptor when they engage with HLAs^71^. Studies have indicated that featureless epitopes are more likely to select relatively narrow TCR repertoires than epitopes with large exposed features *in vivo*^72^. We next examined the ESA feature of C08-mut9m and C08-mut10m, by employing the PDBePISA server. Of note, ESAs of two different T cell epitopes were also calculated as benchmarks. One is HLA-A2–M1, a viral antigen “M1” (M1_58-66_ from the IAV) in complex with HLA-A*02:01(A2-M1, in Fig. 4a), which is considered a featureless epitope^73^. In contrast, the viral epitope HLA-A2-RT is a reverse transcriptase peptide (RT_468-476_ from HIV) in complex with HLA-A*02:01(called A2-RT, in Fig. 4a), and is considered as a largely exposed epitope^74^. After calculation, we found that mut9m has the smallest ESA at 240 Å^2^, even less than the well-known featureless M1 peptide (251 Å^2^, in Fig. 4a-c). However, mut10m has a relatively large ESA at 317 Å^2^, which is comparable to the typical largely exposed antigen RT (330 Å^2^, in Fig. 4a-c). These data suggested that mut9m provides relatively less ESA than canonical T cell antigens. We thus postulated that the diversity of C08-mut9m-restricted TCRs may be constrained *in vivo* because of the featureless area available for TCR recognition.

**Figure 4.**
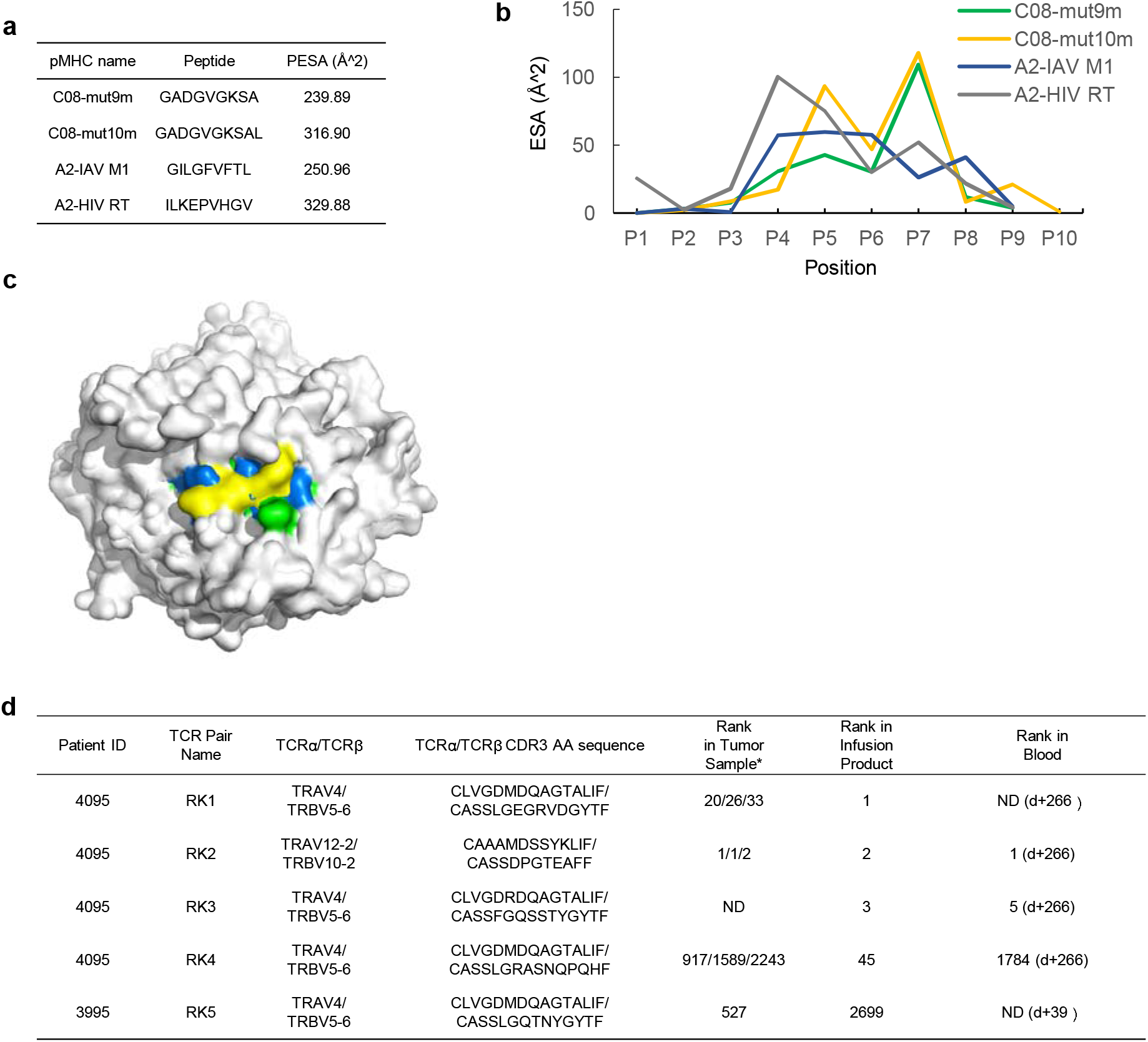
Different peptide exposed surface area (ESA) between C08-mut9m and C08-mut10m. **a**, Peptide exposed surface area (ESA) of four pMHCs. ESA of A2-IAV M1 complex was calculated based on 2VLL (PDB ID). ESA of A2-HIV RT complex was calculated based on 2×4U. **b**, ESA of individual residues at each position of four peptides within HLAs. **c**, Mut10m exhibits relatively large ESA compared to A2-IAV M1 (the featureless benchmark), C08-mut9m, and C08-mut10m. The HLA backbone surface was shown in grey. IAV M1 (blue), mut9m (green), and mut10m (yellow) peptide surfaces were shown in different colors. **d**, Different T cell fates of KRAS G12D neoantigen restricted T cells reported by previous clinical studies. Rank in tumor sample represents the rank of restricted T cells in patient’s TILs before cell therapy. Rank in infusion products represents the rank of restricted T cells in cell transfer products before cell transfer. Rank in blood represents the rank of restricted T cells in patients’ peripheral blood after cell transfer (d+ represents the day after cell transfer). The rank of T cells with the same TCR pair was validated in one or three different metastatic tumor fragments from the patient. *, tested in three fragments; †, tested in one fragment; ND, not detected.

Studies have suggested that only narrow TCR repertoires can recognize featureless epitope, because of the lack of TCR recognition modes^72,73,75^. To observe the diversity of KRAS G12D neoantigen-specific TCRs in clinical cases, we examined the TCR sequences of those restricted T cell repertoires targeting C08-mut9m (Fig. 4d, patients 3995 and 4095, both expressing HLA-C*08:02)^12,57^. Patient 3995 received ACT (adoptive cell transfer) treatments and did not respond. In this trial, the transferred cell products contain the RK5 (RK herein refers to ‘Restricted KRAS G12D mutation’) T cell repertoire which can recognize C08-mut9m. Patient 4095 received an ACT treatment and observed objective tumor regressions. The RK1, RK3, RK4 T cell clones from this patient were verified to recognize C08-mut9m while the RK2 clone recognized C08-mut10m. All these four T cell clones (RK1, RK3, RK4, RK5) with C08-mut9m restriction were identified to have biased usage of a public TCR pair (TRAV4/TRBV5-1) across two patients. The length and sequence of these TCRα chains were highly restricted, with the same TCR-V region and the “CLVGDxDQAGTALIF” CDR3α motif among the four TCRs (Fig. 4d). TCRβ chains were also restricted at TCR-V region but showed differences at CDR3β region. Generally, the CDR1 and CDR2 loops of TCR can recognize the two conservative α-helixes on the MHC, whereas the CDR3 loops mainly interact with the exposed peptide. Moreover, CDR3β had proved to be the main factor that determines TCR recognition (compared with CDR3α) in many cases, due to greater sequence diversity and extensive contacts to the peptide region^75^. However, in this case, the C08-mut9m restricted TCR sequences of CDR1, CRR2, and CDR3α were found to be consistent across different patients. We thus speculated that these public regions of CDR1, CDR2, and CDR3α are important in the C08-mut9m recognition, rather than the CDR3β regions. In contrast, the C08-mut10m neoepitope did not observe dominant public TCRs in patient 4095 or across different individuals. Collectively, these data suggested that the featureless mut9m neoepitope can be recognized by T cells with public and limited TCRs across different patients.

Previous reports have shown that constrained TCR repertoires are associated with poor control of viral infection^76,77^. However, the correlation between TCR bias and clinical outcome in cancer treatment is unclear. In an exploratory analysis, we examined the TCR bias and clinical performance in the above cases. The C08-mut9m restricted T cell clones with public TCR usage (RK1, RK3 RK4, and RK5) were not the top ranked clonotypes among the TILs (Fig. 4d). However, the RK2 T cells with C08-mut10m restriction showed dominant persistence in both TIL and the blood after cell transfer. While we did not observe a direct correlation between TCR diversity and clinical outcome due to limitations in these clinical data, we still observed short-lived T cell persistence in the presence of the featureless C08-mut9m neoepitope. We thus postulated that cancer patients who receive neoantigen-based immunotherapies with characteristic (relatively large ESA) neoantigens might obtain neoantigen-restricted T cells with better persistence *in vivo*.

## Discussion

How T cells recognize neoantigens as “non-self” is an important question in cancer immunotherapy. In contrast to conventional pathogenic peptides that are completely foreign, neoantigens can differ from self by only a single amino acid. Understanding the factors and mechanisms that contribute to the immunogenicity of neoantigens may instruct the design of future immunotherapies.

Efforts have been made to understand the complexity of neoantigen immunogenicity with a major focus thus far on peptide-MHC binding. Different studies have led to different conclusions^78^. Fritsch *et al*. and Yadav *et al*. suggest that immunogenic neoantigens are more commonly mutated at a TCR-contacting residue, while Duan *et al*. suggested that neoantigen substitutions at MHC anchor residues may be more immunogenic^7,18,28^. Further, using mouse models, Capietto *et al*. showed that increased affinity relative to the corresponding wild-type peptide can influence neoantigen immunogenicity^29^. We found that mutations occur more frequently at TCR-contacting positions rather than MHC-contacting positions in the immunogenic dataset. However, amino acid biochemical properties might not be critical for neoantigen filtering for candidates with TCR-contacting position mutation. For mutations that occur at the MHC anchor positions, we showed that the NP rule of anchor mutated neoantigens exists pervasively. Also, we found that the NP rule combines with the MHC binding predictor NetMHCpan 4.0 to enhance the prioritization of neoantigen candidates. In a sense, the NP feature could be captured by existing neoantigen predictors through the comparison of binding differences between wild-type and mutant peptides^18,79,80^. However, the NP rule we provide here is a binary feature that can provide a direct understanding of neoantigen immunogenicity. Nevertheless, the robustness of the NP rule will need to be tested with larger sample sizes.

We also performed structural studies to obtain insights into the NP rule. We found that anchor mutated neoantigens that followed the NP rule can generate new surfaces and features for T cell recognition that the wild-type peptide cannot. In contrast, the anchor mutated neopeptide from DPAGT1 with PP rule exhibited low immunogenicity in clinical treatment. It is possible that most of the neoepitope restricted T cells were also cross-reactive with wild-type and thus were removed by negative selection. Our data also suggests that neoantigen exposed surface area (ESA) might be a factor that influences TCR diversity and clinical outcome, but more experimental data and neoantigen-MHC structures are needed to fully understand the relevance between ESA and TCR diversity.

Our study showed three possible neoantigen binding models within the context of MHC (Supplementary Fig. 5). Model A represents the situation in which a mutation occurs in the peptide that contacts TCR. Model B represents the situation in which a mutation occurs at a non-anchor MHC-contacting region and therefore might be least immunogenic. Model C represents the situation in which a mutation occurs at MHC anchor position. Anchor mutations may not change the TCR-contacting surface but instead lead to enhanced de novo presentation of the peptide for TCR recognition.

KRAS G12D is one of the most common driver mutations that leads to oncogenesis^81^. It is also indicative of poor prognosis with poor response to standard cancer treatments. The transfer of HLA-C*08:02-restricted T cells targeting KRAS G12D neoantigens was associated with clinical response. Based on these pMHC structures, further research could be undertaken to increase the immunogenicity and stability of the KRAS G12D-C*08:02 neoepitope, by making modifications of agonist peptides or by screening non-natural synthetic epitopes^82,83^. Our structures could also be used to design artificial receptors/proteins that bind mutant KRAS peptides based on synthetic biology approaches.

## Acknowledgments

We thank the staff from BL17U1/BL18U1/BL19U1 beamline of National Center for Protein Sciences Shanghai (NCPSS) at Shanghai Synchrotron Radiation Facility, for assistance during crystal data collection. We would like to thank Chang-Yi Ma (The Chinese University of Hong Kong, China) for the advice on data analysis. This research was funded by the National Institutes of Health Grants AI018785 and AI135374 (to P.M.); the National Natural Science Foundation of China 31870728 and 31470738 (to L.Y.); the National Basic Research Program of China 2014CB910103 (to L.Y.); the Science Foundation of Wuhan University 2042016kf0169 (to L.Y.).

## Author contributions

P.B., L.Y., and P.M. designed all the experiments; P.B. performed all experiments; P.B. and J.X. collected neoantigen data; P.B., L.Y., P.M., J.W.K., M.W., Y.L., Y.Z., S.K.C., and J.X. analyzed neoantigen data; P.B., Q.Z., P.W. and H.D. determined crystal structures; P.B., L.Y., and P.W. analyzed crystal structures; P.B., L.Y., P.M., E.T., J.W.K., and S.K.C. wrote the manuscript.

## Competing interests

The authors declare that they have no conflict of interest.

## Supplementary Figure Legends

**Supplementary Figure 1.**
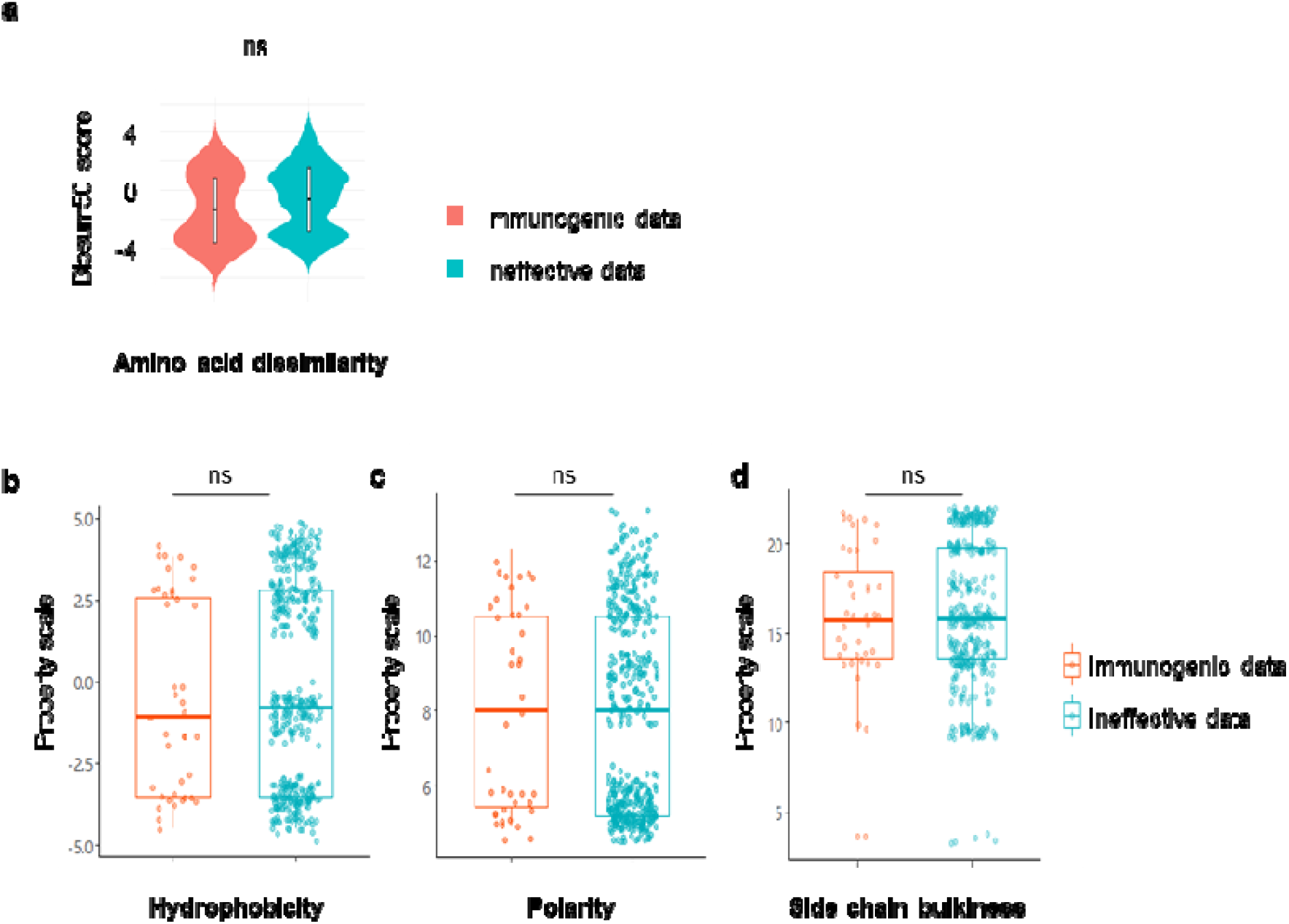
Comparison of four amino acid biochemical properties between immunogenic and ineffective neoantigen data harboring TCR-contacting position mutation. **a**, Comparison of amino acid dissimilarity of neoantigen pairs based on immunogenic and ineffective data. Normalized score was calculated based on the BLOSUM50 matrix (p= 0.0671, n=40 (immunogenic); n=426 (ineffective), Wilcoxon rank sum; ns, not significant). **b**, Comparison of mutation hydrophobicity based on immunogenic and ineffective data. Hydrophobicity property scales for amino acids was recorded in the “amino acid property scales” table (p= 0.4039, n=40 (immunogenic); n=426 (ineffective), Wilcoxon rank sum; ns, not significant). **c**, Comparison of mutation polarity based on immunogenic and ineffective data. Polarity property scales for each amino acid were recorded in the “amino acid property scales” table (p= 0.5121, n=40 (immunogenic); n=426 (ineffective), Wilcoxon rank sum; ns, not significant). **d**, Comparison of mutation side chain bulkiness based on immunogenic and ineffective data. Side chain bulkiness scales for amino acids were recorded in the “amino acid property scales” table (p= 0.8995, n=40 (immunogenic); n=426 (ineffective), Wilcoxon rank sum; ns, not significant).

**Supplementary Figure 2.**
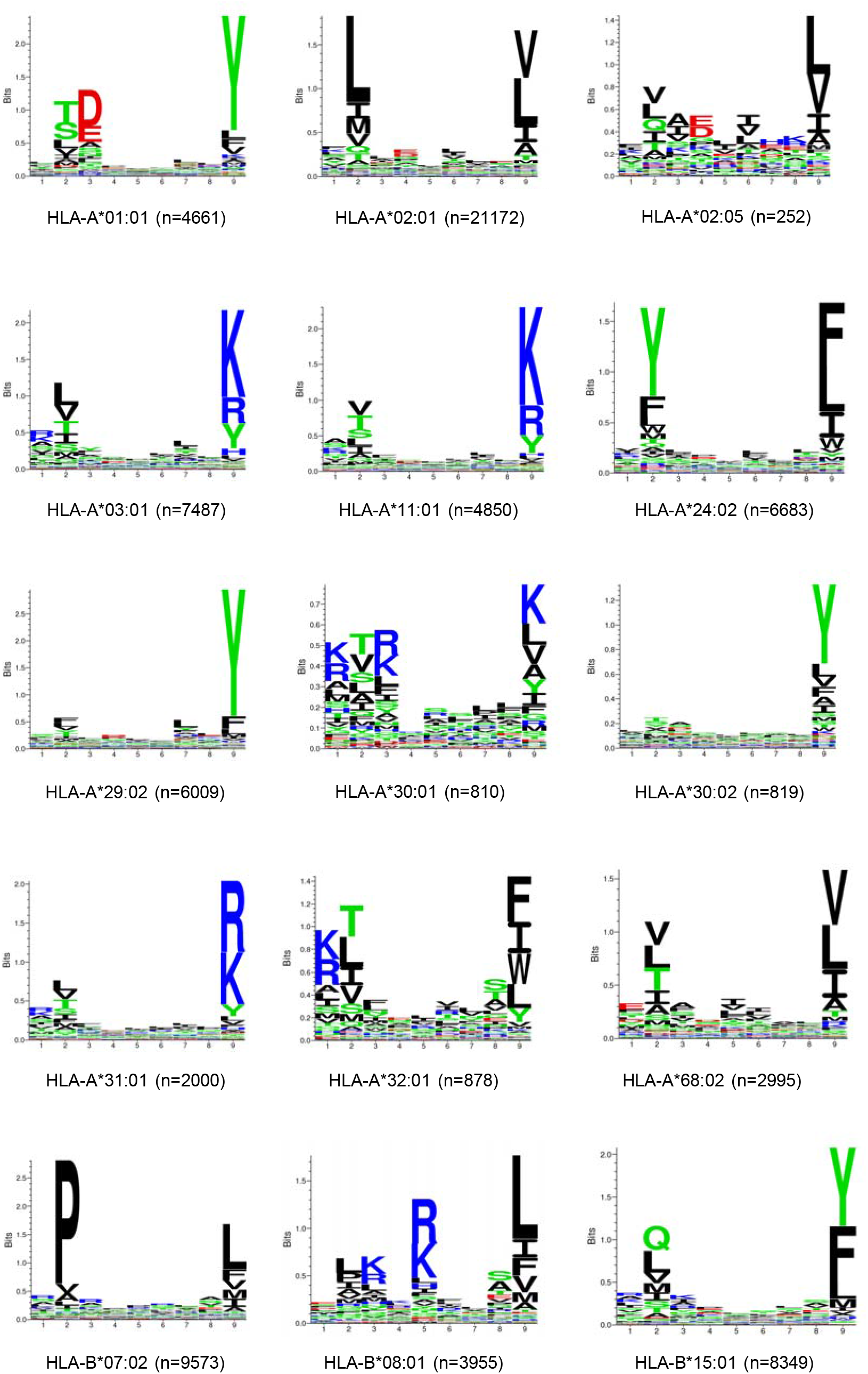
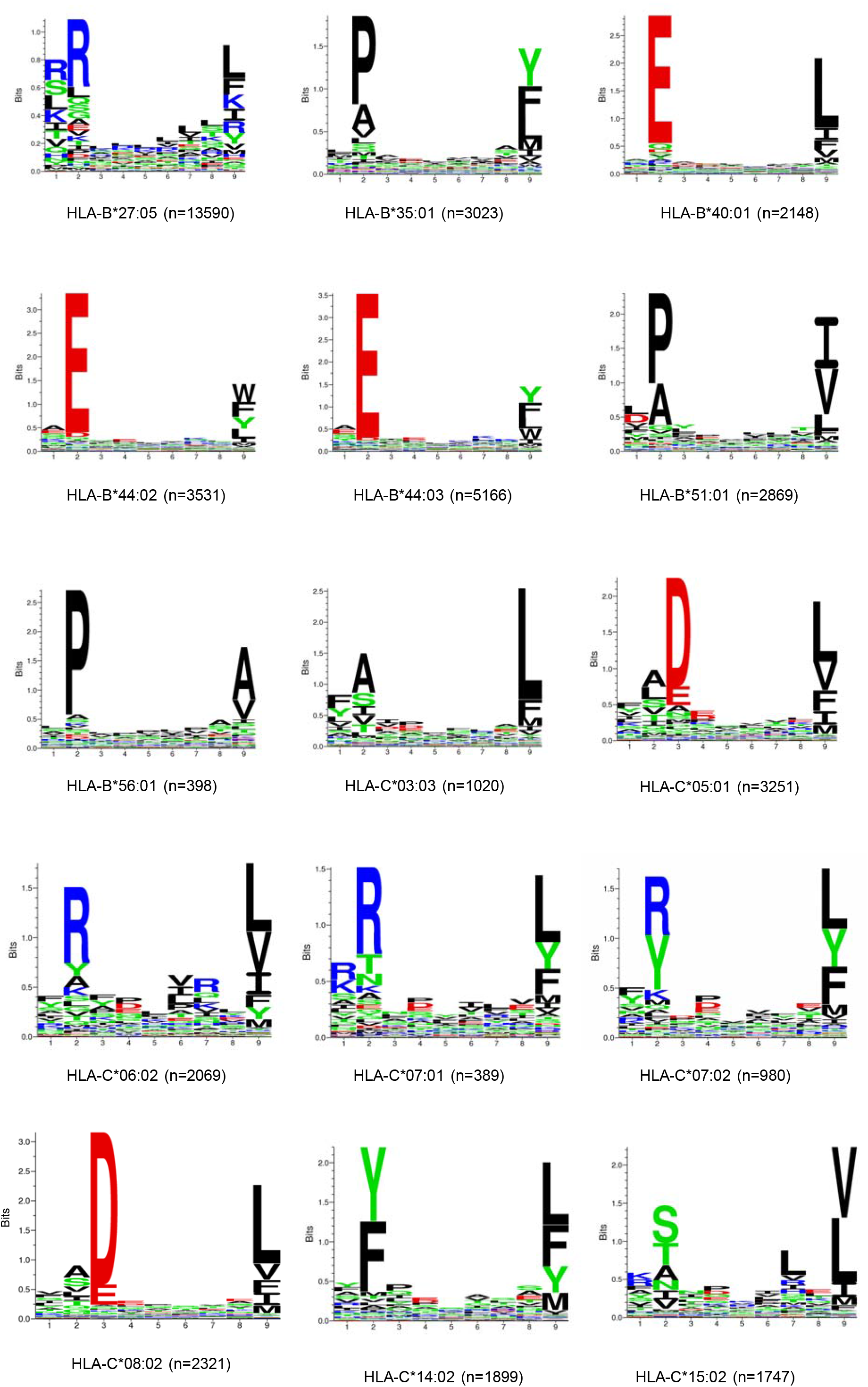
Binding profiles of nonameric peptides of 30 HLA alleles. Sequence logos of 30 HLA alleles were generated from peptide-binding matrices using the Seq2Logo. Peptide libraries (9mer peptides) were obtained from IEDB. The sample size of each allele was shown in the figure.

**Supplementary Figure 3.**
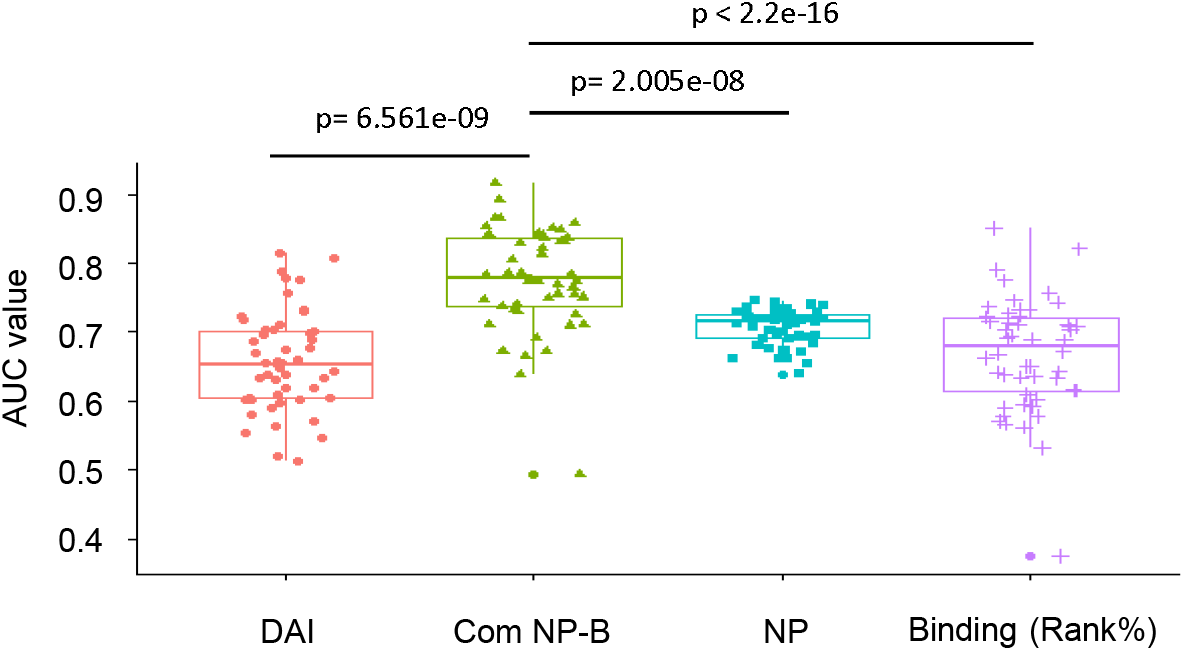
AUC value testing using 50-fold cross-validation. The 50-fold cross-validation (2/3rd random resampling) was performed within the exploration set. Differences of AUC values were determined using paired T-test.

**Supplementary Figure 4.**
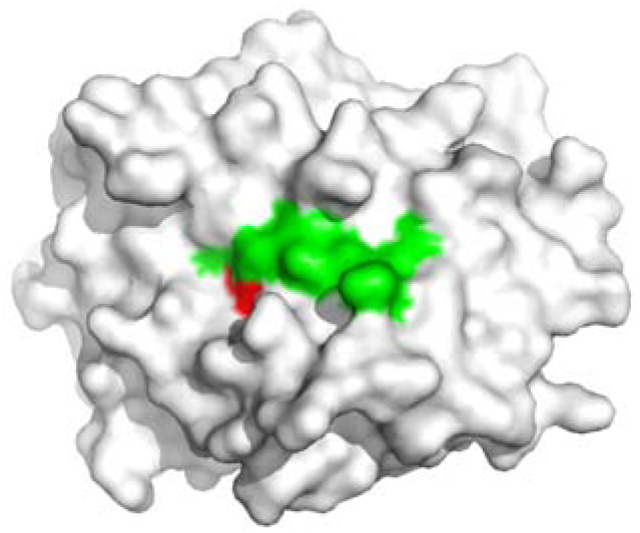
Additional TCR accessible surface of C08-mut9m. P3D of mut9m can provide an additional TCR accessible surface. HLA-C*08:02 (grey), 9mer peptide (green), and the P3D side chain (red) were showed in different colors.

**Supplementary Figure 5.**
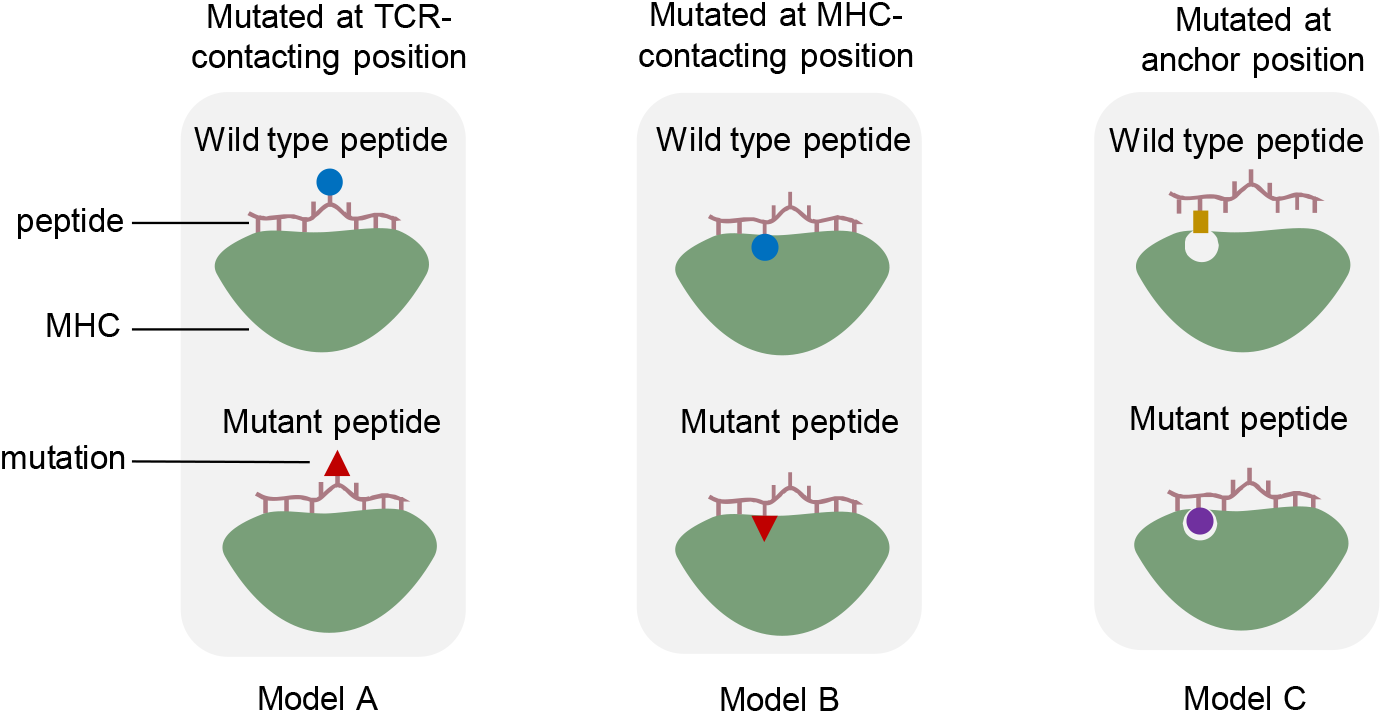
The topology of different neoantigen-MHC binding models. Model A shows the neoantigens with mutations at TCR-contacting positions. The blue circle represents wild-type residues. The red triangle represents the mutant residues. Model B shows the neoantigens with mutations at MHC-contacting positions. The blue circle represents wild-type residues. Model C shows the peptides which have mutations at anchor positions. The yellow rectangle represents wild-type residues that cannot fit the binding pocket on MHC. The purple circle represents mutated residues which are preferentially selected by MHC.

**Supplementary Table 1. Immunogenic Neopeptide Dataset (IND)**

**Supplementary Table 2. Raw ineffective Neopeptide Dataset (RIEND)**

**Supplementary Table 3. Ineffective Neopeptide Dataset (IEND)**

(**see in separated EXCEL files)**

**Supplementary Table 4.** Amino acid property scales

|  | Hydrophobicity | Polarity | Bulkiness |
| --- | --- | --- | --- |
| Alanine/A | 1.8 | 8 | 11.5 |
| Cysteine/C | 2.5 | 5.5 | 13.46 |
| Aspartic acid/D | -3.5 | 13 | 11.68 |
| Glutamic acid/E | -3.5 | 12.3 | 13.57 |
| Phenylalanine/F | 2.8 | 5.2 | 19.8 |
| Glycine/G | -0.4 | 9 | 3.4 |
| Histidine/H | -3.2 | 10.4 | 13.69 |
| Isoleucine/I | 4.5 | 5.2 | 21.4 |
| Lysine/K | -3.9 | 11.3 | 15.71 |
| Leucine/L | 3.8 | 4.9 | 21.4 |
| Methionine/M | 1.9 | 5.7 | 16.25 |
| Asparagine/N | -3.5 | 11.6 | 12.82 |
| Proline/P | -1.6 | 8 | 17.43 |
| Glutamine/Q | -3.5 | 10.5 | 14.45 |
| Arginine/R | -4.5 | 10.5 | 14.28 |
| Serine/S | -0.8 | 9.2 | 9.47 |
| Threonine/T | -0.7 | 8.6 | 15.77 |
| Valine/V | 4.2 | 5.9 | 21.57 |
| Tryptophan/W | -0.9 | 5.4 | 21.67 |
| Tyrosine/Y | -1.3 | 6.2 | 18.03 |

Independent numeric scales for different amino acid biochemical properties (including amino acid "hydrophobicity", "polarity" and "bulkiness". This table was described by *Chowell et al.* (PNAS, 2015, E1754–E1762).

**Supplementary Table 5.** Classification of anchor mutated neoantigens based on MHC preference

|  | N->N<br>frequency | N->P<br>frequency | P->N<br>frequency | P->P<br>frequency | Number of<br>peptide |
| --- | --- | --- | --- | --- | --- |
| **Anchor mutated neoantigens (immunogenic)** | 3.7% (1) | 96.3% (26) | 0% (0) | 0% (0) | 27 |
| **Anchor mutated neopeptides (ineffective)** | 27.76% (118) | 56% (238) | 6.82% (29) | 9.41% (40) | 425 |

Data come from Immunogenic Neoantigen Dataset (IND) and Ineffective Neopeptide Dataset (IEND). Peptides defined with anchor position were calculated in this analysis.

**Supplementary Table 6.** Predicted binding scores and biophysical thermal stability of KRAS peptides in the context of HLA-C*08:02

| Peptide | pRank (%) | Thermal stability (T<sub>m</sub>, °C ) |
| --- | --- | --- |
| GADGVGKSA (mut9m) | 1.5435 (weak binder) | 50±1.6 |
| GADGVGKSAL (mut10m) | 0.3508 (strong binder) | 48±1.1 |
| GAGGVGKSA (wt9m) | 15.00 (non-binder) | N |
| GAGGVGKSAL (wt10m) | 8.0828 (non-binder) | N |

All tests were performed in the presence of HLA-C\*08:02. Peptide binding prediction was performed by NetMHCpan 4.0 server. The percentile rank (pRank, %) represents the predicted affinity compared to a set of random natural peptides. Strong binders are defined as having %rank<0.5, and weak binders with %rank<2. Peptides that have %rank>2 can be treated as non-binders.
The T<sub>m</sub>, or thermal melt point, represents the temperature for which 50% of the protein is unfolded. The wt9m and wt10mer peptides failed to refold with HLA-C\*08:02 and thus did not test the thermal stability further. N, did not test.

**Supplementary Table 7.** Data collection and refinement statistics (molecular replacement).

|  | C08-mut9m | C08-mut10m | Kb-8mV | Kb-8mL |
| --- | --- | --- | --- | --- |
| Resolution range | 42.55-1.9 (1.968-1.9) | 51.13-1.9 (1.968 -1.9) | 57.76- 2.4 (2.486-2.4) | 68.67-2.5 (2.589-2.5) |
| Space group | P 21 21 21 | P 21 21 21 | C 1 2 1 | C 1 2 1 |
| Unit cell | 68.3 75.9 78.0<br>90 90 90 | 68.9 76.2 78.7<br>90 90 90 | 107.2 69.4 70.4<br>90 103.8 90 | 106.6 68.0 70.7<br>90 103.9 90 |
| Total reflections | 64768 (6323) | 66521 (6583) | 37661 (3734) | 33008 (3336) |
| Unique reflections | 32450 (3174) | 33289 (3294) | 19013 (1886) | 16646 (1679) |
| Multiplicity | 2.0 (2.0) | 2.0 (2.0) | 2.0 (2.0) | 2.0 (2.0) |
| Completeness (%) | 0.96 (0.99) | 1.00 (1.00) | 0.96 (0.96) | 0.97 (0.99) |
| Mean I/sigma(I) | 7.15 (4.02) | 9.26 (5.12) | 14.38 (4.80) | 11.60 (2.85) |
| Wilson B-factor (Å<sup>2</sup>) | 10.39 | 20.99 | 38.48 | 46.21 |
| R-merge | 0.04092 (0.1168) | 0.03331 (0.1176) | 0.02281 (0.1159) | 0.02767 (0.1982) |
| R-meas | 0.05787 (0.1652) | 0.04711 (0.1663) | 0.03225 (0.1639) | 0.03913 (0.2803) |
| CC1/2 | 0.995 (0.963) | 0.996 (0.956) | 0.999 (0.979) | 0.999 (0.944) |
| CC* | 0.999 (0.99) | 0.999 (0.989) | 1 (0.995) | 1 (0.985) |
| Reflections used in refinement | 31350 (2603) | 33285 (3294) | 18981 (1879) | 16622 (1674) |
| Reflections used for R-free | 1936 (155) | 2001 (198) | 1899 (188) | 1663 (167) |
| R-work | 0.2406 | 0.174 | 0.2194 | 0.2153 |
| R-free | 0.2872 | 0.2044 | 0.263 | 0.2934 |
| CC(work) | 0.933 | 0.96 | 0.943 | 0.958 |
| CC(free) | 0.897 | 0.946 | 0.896 | 0.902 |
| Number of non-hydrogen atoms | 3641 | 3542 | 3152 | 3093 |
| macromolecules | 3127 | 3139 | 3044 | 3045 |
| Protein residues | 381 | 382 | 373 | 373 |
| RMS(bonds) (Å) | 0.008 | 0.007 | 0.01 | 0.01 |
| RMS(angles) (°) | 0.93 | 0.93 | 1.08 | 1.19 |
| Ramachandran favored (%) | 96 | 99 | 94 | 94 |
| Ramachandran allowed (%) | 3.2 | 1.3 | 5.3 | 5.1 |
| Ramachandran outliers (%) | 0.53 | 0 | 0.28 | 0.85 |
| Rotamer outliers (%) | 2.1 | 1.8 | 4.3 | 2.8 |
| Clashscore | 8.39 | 4.91 | 18.38 | 15.2 |
| Average B-factor (Å<sup>2</sup>) | 17.27 | 25.67 | 50.83 | 59.08 |
| Macromolecules | 16.15 | 24.62 | 50.95 | 59.2 |
| solvent | 24.05 | 33.81 | 47.71 | 51.52 |

Statistics for the highest-resolution shell are shown in parentheses.

## Notes

### Competing Interest Statement

The authors have declared no competing interest.

### Summary of Updates

Published edition revision.

## References

1. Vesely MD, Kershaw MH, Schreiber RD, Smyth MJ. Natural Innate and Adaptive Immunity to Cancer. Annu. Rev. Immunol. 2011;29(1):235–271. doi:10.1146/annurev-immunol-031210-101324

2. Sykulev Y, Joo M, Vturina I, Tsomides TJ, Eisen HN. Evidence that a Single Peptide–MHC Complex on a Target Cell Can Elicit a Cytolytic T Cell Response. Immunity. 1996;4(6):565–571. doi:10.1016/S1074-7613(00)80483-5

3. Tumeh PC, Harview CL, Yearley JH, Shintaku IP, Taylor EJM, Robert L, Chmielowski B, Spasic M, Henry G, Ciobanu V, et al. PD-1 blockade induces responses by inhibiting adaptive immune resistance. Nature. 2014;515(7528):568–571. doi:10.1038/nature13954

4. Schumacher TN, Schreiber RD. Neoantigens in cancer immunotherapy. Science. 2015;348(6230):69–74. doi:10.1126/science.aaa4971

5. Rizvi NA, Hellmann MD, Snyder A, Kvistborg P, Makarov V, Havel JJ, Lee W, Yuan J, Wong P, Ho TS, et al. Mutational landscape determines sensitivity to PD-1 blockade in non-small cell lung cancer. Science. 2015;348(6230):124–128. doi:10.1126/science.aaa1348

6. Wang R-F, Wang HY. Immune targets and neoantigens for cancer immunotherapy and precision medicine. Cell Res. 2017;27(1):11–37. doi:10.1038/cr.2016.155

7. Yadav M, Jhunjhunwala S, Phung QT, Lupardus P, Tanguay J, Bumbaca S, Franci C, Cheung TK, Fritsche J, Weinschenk T, et al. Predicting immunogenic tumour mutations by combining mass spectrometry and exome sequencing. Nature. 2014;515(7528):572–576. doi:10.1038/nature14001

8. Coulie PG, Van den Eynde BJ, van der Bruggen P, Boon T. Tumour antigens recognized by T lymphocytes: at the core of cancer immunotherapy. Nat. Rev. Cancer. 2014;14(2):135–146. doi:10.1038/nrc3670

9. Gascoigne NRJ, Rybakin V, Acuto O, Brzostek J. TCR Signal Strength and T Cell Development. Annu. Rev. Cell Dev. Biol. 2016;32(1):327–348. doi:10.1146/annurev-cellbio-111315-125324

10. Matsushita H, Vesely MD, Koboldt DC, Rickert CG, Uppaluri R, Magrini VJ, Arthur CD, White JM, Chen Y-S, Shea LK, et al. Cancer exome analysis reveals a T-cell-dependent mechanism of cancer immunoediting. Nature. 2012;482(7385):400–404. doi:10.1038/nature10755

11. Robbins PF, Lu Y-C, El-Gamil M, Li YF, Gross C, Gartner J, Lin JC, Teer JK, Cliften P, Tycksen E, et al. Mining exomic sequencing data to identify mutated antigens recognized by adoptively transferred tumor-reactive T cells. Nat. Med. 2013;19(6):747–752. doi:10.1038/nm.3161

12. Tran E, Robbins PF, Lu Y-C, Prickett TD, Gartner JJ, Jia L, Pasetto A, Zheng Z, Ray S, Groh EM, et al. T-Cell Transfer Therapy Targeting Mutant KRAS in Cancer. N. Engl. J. Med. 2016;375(23):2255– 2262. doi:10.1056/NEJMoa1609279

13. Ott PA, Hu Z, Keskin DB, Shukla SA, Sun J, Bozym DJ, Zhang W, Luoma A, Giobbie-Hurder A, Peter L, et al. An immunogenic personal neoantigen vaccine for patients with melanoma. Nature. 2017;547(7662):217–221. doi:10.1038/nature22991

14. Sahin U, Derhovanessian E, Miller M, Kloke B-P, Simon P, Löwer M, Bukur V, Tadmor AD, Luxemburger U, Schrörs B, et al. Personalized RNA mutanome vaccines mobilize poly-specific therapeutic immunity against cancer. Nature. 2017;547(7662):222–226. doi:10.1038/nature23003

15. Melief CJM. Precision T-cell therapy targets tumours. Nature. 2017;547(7662):165–167. doi:10.1038/nature23093

16. Chen F, Zou Z, Du J, Su S, Shao J, Meng F, Yang J, Xu Q, Ding N, Yang Y, et al. Neoantigen identification strategies enable personalized immunotherapy in refractory solid tumors. J. Clin. Invest. 2019;129(5):2056–2070. doi:10.1172/JCI99538

17. van Rooij N, van Buuren MM, Philips D, Velds A, Toebes M, Heemskerk B, van Dijk LJA, Behjati S, Hilkmann H, el Atmioui D, et al. Tumor Exome Analysis Reveals Neoantigen-Specific T-Cell Reactivity in an Ipilimumab-Responsive Melanoma. J. Clin. Oncol. 2013;31(32):e439–e442. doi:10.1200/JCO.2012.47.7521

18. Duan F, Duitama J, Al Seesi S, Ayres CM, Corcelli SA, Pawashe AP, Blanchard T, McMahon D, Sidney J, Sette A, et al. Genomic and bioinformatic profiling of mutational neoepitopes reveals new rules to predict anticancer immunogenicity. J. Exp. Med. 2014;211(11):2231–2248. doi:10.1084/jem.20141308

19. Cohen CJ, Gartner JJ, Horovitz-Fried M, Shamalov K, Trebska-McGowan K, Bliskovsky VV, Parkhurst MR, Ankri C, Prickett ToddD, Crystal JS, et al. Isolation of neoantigen-specific T cells from tumor and peripheral lymphocytes. J. Clin. Invest. 2015;125(10):3981–3991. doi:10.1172/JCI82416

20. Carreno BM, Magrini V, Becker-Hapak M, Kaabinejadian S, Hundal J, Petti AA, Ly A, Lie W-R, Hildebrand WH, Mardis ER, et al. A dendritic cell vaccine increases the breadth and diversity of melanoma neoantigen-specific T cells. Science. 2015;348(6236):803. doi:10.1126/science.aaa3828

21. Stronen E, Toebes M, Kelderman S, van Buuren MM, Yang W, van Rooij N, Donia M, Boschen M-L, Lund-Johansen F, Olweus J, et al. Targeting of cancer neoantigens with donor-derived T cell receptor repertoires. Science. 2016;352(6291):1337–1341. doi:10.1126/science.aaf2288

22. Anagnostou V, Smith KN, Forde PM, Niknafs N, Bhattacharya R, White J, Zhang T, Adleff V, Phallen J, Wali N, et al. Evolution of Neoantigen Landscape during Immune Checkpoint Blockade in Non–Small Cell Lung Cancer. Cancer Discov. 2017;7(3):264–276. doi:10.1158/2159-8290.CD-16-0828

23. Zacharakis N, Chinnasamy H, Black M, Xu H, Lu Y-C, Zheng Z, Pasetto A, Langhan M, Shelton T, Prickett T, et al. Immune recognition of somatic mutations leading to complete durable regression in metastatic breast cancer. Nat. Med. 2018;24(6):724–730. doi:10.1038/s41591-018-0040-8

24. Jurtz VI, Paul S, Andreatta M, Marcatili P, Peters B, Nielsen M. NetMHCpan 4.0: Improved peptide-MHC class I interaction predictions integrating eluted ligand and peptide binding affinity data. 2017. doi:10.1101/149518

25. Andreatta M, Nielsen M. Gapped sequence alignment using artificial neural networks: application to the MHC class I system. Bioinformatics. 2016;32(4):511–517. doi:10.1093/bioinformatics/btv639

26. Vitiello A, Zanetti M. Neoantigen prediction and the need for validation. Nat. Biotechnol. 2017;35(9):815–817. doi:10.1038/nbt.3932

27. Bulik-Sullivan B, Busby J, Palmer CD, Davis MJ, Murphy T, Clark A, Busby M, Duke F, Yang A, Young L, et al. Deep learning using tumor HLA peptide mass spectrometry datasets improves neoantigen identification. Nat. Biotechnol. 2019;37(1):55–63. doi:10.1038/nbt.4313

28. Fritsch EF, Rajasagi M, Ott PA, Brusic V, Hacohen N, Wu CJ. HLA-Binding Properties of Tumor Neoepitopes in Humans. Cancer Immunol. Res. 2014;2(6):522–529. doi:10.1158/2326-6066.CIR-13-0227

29. Capietto A-H, Jhunjhunwala S, Pollock SB, Lupardus P, Wong J, Hänsch L, Cevallos J, Chestnut Y, Fernandez A, Lounsbury N, et al. Mutation position is an important determinant for predicting cancer neoantigens. J. Exp. Med. 2020;217(4). doi:10.1084/jem.20190179

30. Henikoff S, Henikoff JG. Amino acid substitution matrices from protein blocks. Proc. Natl. Acad. Sci. 1992;89(22):10915–10919. doi:10.1073/pnas.89.22.10915

31. Chowell D, Krishna S, Becker PD, Cocita C, Shu J, Tan X, Greenberg PD, Klavinskis LS, Blattman JN, Anderson KS. TCR contact residue hydrophobicity is a hallmark of immunogenic CD8 ^+^ T cell epitopes. Proc. Natl. Acad. Sci. 2015;112(14):E1754–E1762. doi:10.1073/pnas.1500973112

32. Ghorani E, Rosenthal R, McGranahan N, Reading JL, Lynch M, Peggs KS, Swanton C, Quezada SA. Differential binding affinity of mutated peptides for MHC class I is a predictor of survival in advanced lung cancer and melanoma. Ann. Oncol. 2018;29(1):271–279. doi:10.1093/annonc/mdx687

33. Rammensee H-G, Bachmann J, Emmerich NPN, Bachor OA, Stevanović S. SYFPEITHI: database for MHC ligands and peptide motifs. Immunogenetics. 1999;50(3–4):213–219. doi:10.1007/s002510050595

34. Vita R, Mahajan S, Overton JA, Dhanda SK, Martini S, Cantrell JR, Wheeler DK, Sette A, Peters B. The Immune Epitope Database (IEDB): 2018 update. Nucleic Acids Res. 2019;47(D1):D339–D343. doi:10.1093/nar/gky1006

35. Thomsen MCF, Nielsen M. Seq2Logo: a method for construction and visualization of amino acid binding motifs and sequence profiles including sequence weighting, pseudo counts and two-sided representation of amino acid enrichment and depletion. Nucleic Acids Res. 2012;40(W1):W281– W287. doi:10.1093/nar/gks469

36. Yin L, Huseby E, Scott-Browne J, Rubtsova K, Pinilla C, Crawford F, Marrack P, Dai S, Kappler JW. A Single T Cell Receptor Bound to Major Histocompatibility Complex Class I and Class II Glycoproteins Reveals Switchable TCR Conformers. Immunity. 2011;35(1):23–33. doi:10.1016/j.immuni.2011.04.017

37. Macdonald WA, Chen Z, Gras S, Archbold JK, Tynan FE, Clements CS, Bharadwaj M, Kjer-Nielsen L, Saunders PM, Wilce MCJ, et al. T Cell Allorecognition via Molecular Mimicry. Immunity. 2009;31(6):897–908. doi:10.1016/j.immuni.2009.09.025

38. Battye TGG, Kontogiannis L, Johnson O, Powell HR, Leslie AGW. iMOSFLM?: a new graphical interface for diffraction-image processing with MOSFLM. Acta Crystallogr. D Biol. Crystallogr. 2011;67(4):271–281. doi:10.1107/S0907444910048675

39. Winn MD, Ballard CC, Cowtan KD, Dodson EJ, Emsley P, Evans PR, Keegan RM, Krissinel EB, Leslie AGW, McCoy A, et al. Overview of the CCP 4 suite and current developments. Acta Crystallogr. D Biol. Crystallogr. 2011;67(4):235–242. doi:10.1107/S0907444910045749

40. McCoy AJ, Grosse-Kunstleve RW, Adams PD, Winn MD, Storoni LC, Read RJ. Phaser crystallographic software. J. Appl. Crystallogr. 2007;40(4):658–674. doi:10.1107/S0021889807021206

41. Emsley P, Cowtan K. Coot?: model-building tools for molecular graphics. Acta Crystallogr. D Biol. Crystallogr. 2004;60(12):2126–2132. doi:10.1107/S0907444904019158

42. Adams PD, Grosse-Kunstleve RW, Hung L-W, Ioerger TR, McCoy AJ, Moriarty NW, Read RJ, Sacchettini JC, Sauter NK, Terwilliger TC. PHENIX?: building new software for automated crystallographic structure determination. Acta Crystallogr. D Biol. Crystallogr. 2002;58(11):1948– 1954. doi:10.1107/S0907444902016657

43. Hogan KT, Eisinger DP, Cupp SB, Lekstrom KJ, Deacon DD, Shabanowitz J, Hunt DF, Engelhard VH, Slingluff CL, Ross MM. The Peptide Recognized by HLA-A68.2-restricted, Squamous Cell Carcinoma of the Lung-specific Cytotoxic T Lymphocytes Is Derived from a Mutated <em>Elongation Factor 2</em> Gene. Cancer Res. 1998;58(22):5144.

44. McGranahan N, Furness AJS, Rosenthal R, Ramskov S, Lyngaa R, Saini SK, Jamal-Hanjani M, Wilson GA, Birkbak NJ, Hiley CT, et al. Clonal neoantigens elicit T cell immunoreactivity and sensitivity to immune checkpoint blockade. Science. 2016;351(6280):1463–1469.doi:10.1126/science.aaf1490

45. Robbins PF. A mutated beta-catenin gene encodes a melanoma-specific antigen recognized by tumor infiltrating lymphocytes. J. Exp. Med. 1996;183(3):1185–1192. doi:10.1084/jem.183.3.1185

46. Zorn E, Hercend T. A natural cytotoxic T cell response in a spontaneously regressing human melanoma targets a neoantigen resulting from a somatic point mutation. Eur. J. Immunol. 1999;29(2):592–601. doi:10.1002/(SICI)1521-4141(199902)29:02<592::AID-IMMU592>3.0.CO;2-2

47. Chiari R, Foury F, De Plaen E, Baurain J-F, Thonnard J, G. Coulie P. Two Antigens Recognized by Autologous Cytolytic T Lymphocytes on a Melanoma Result from a Single Point Mutation in an Essential Housekeeping Gene. Cancer Res. 1999;59(22):5785.

48. Echchakir H, Mami-Chouaib F, Vergnon I, Baurain J-F, Karanikas V, Chouaib S, Coulie PG. A Point Mutation in the α-Actinin-4 Gene Generates an Antigenic Peptide Recognized by Autologous Cytolytic T Lymphocytes on a Human Lung Carcinoma. Cancer Res. 2001;61(10):4078.

49. Guéguen M, Patard J-J, Gaugler B, Brasseur F, Renauld J-C, Van Cangh PJ, Boon T, Van den Eynde BJ. An Antigen Recognized by Autologous CTLs on a Human Bladder Carcinoma. J. Immunol. 1998;160(12):6188.

50. Zhou J, Dudley ME, Rosenberg SA, Robbins PF. Persistence of Multiple Tumor-Specific T-Cell Clones Is Associated with Complete Tumor Regression in a Melanoma Patient Receiving Adoptive Cell Transfer Therapy: J. Immunother. 2005;28(1):53–62. doi:10.1097/00002371-200501000-00007

51. Sensi M, Nicolini G, Zanon M, Colombo C, Molla A, Bersani I, Lupetti R, Parmiani G, Anichini A. Immunogenicity without Immunoselection: A Mutant but Functional Antioxidant Enzyme Retained in a Human Metastatic Melanoma and Targeted by CD8+ T Cells with a Memory Phenotype. Cancer Res. 2005:10.

52. Lennerz V, Fatho M, Gentilini C, Frye RA, Lifke A, Ferel D, Wolfel C, Huber C, Wolfel T. The response of autologous T cells to a human melanoma is dominated by mutated neoantigens. Proc. Natl. Acad. Sci. 2005;102(44):16013–16018. doi:10.1073/pnas.0500090102

53. Takenoyama M, Baurain J-F, Yasuda M, So T, Sugaya M, Hanagiri T, Sugio K, Yasumoto K, Boon T, Coulie PG. A point mutation in the NFYC gene generates an antigenic peptide recognized by autologous cytolytic T lymphocytes on a human squamous cell lung carcinoma: NF-YC Mutated Peptide-Reactive CTL in NSCLC. Int. J. Cancer. 2006;118(8):1992–1997. doi:10.1002/ijc.21594

54. Ito D, Visus C, Hoffmann TK, Balz V, Bier H, Appella E, Whiteside TL, Ferris RL, DeLeo AB. Immunological characterization of missense mutations occurring within cytotoxic T cell-defined p53 epitopes in HLA-A*0201+ squamous cell carcinomas of the head and neck. Int. J. Cancer. 2007;120(12):2618–2624. doi:10.1002/ijc.22584

55. Wick DA, Webb JR, Nielsen JS, Martin SD, Kroeger DR, Milne K, Castellarin M, Twumasi-Boateng K, Watson PH, Holt RA, et al. Surveillance of the Tumor Mutanome by T Cells during Progression from Primary to Recurrent Ovarian Cancer. Clin. Cancer Res. 2014;20(5):1125–1134. doi:10.1158/1078-0432.CCR-13-2147

56. Lu Y-C, Yao X, Crystal JS, Li YF, El-Gamil M, Gross C, Davis L, Dudley ME, Yang JC, Samuels Y, et al. Efficient Identification of Mutated Cancer Antigens Recognized by T Cells Associated with Durable Tumor Regressions. Clin. Cancer Res. 2014;20(13):3401–3410. doi:10.1158/1078-0432.CCR-14-0433

57. Tran E, Ahmadzadeh M, Lu Y-C, Gros A, Turcotte S, Robbins PF, Gartner JJ, Zheng Z, Li YF, Ray S, et al. Immunogenicity of somatic mutations in human gastrointestinal cancers. Science. 2015;350(6266):1387–1390. doi:10.1126/science.aad1253

58. Snyder A, Makarov V, Merghoub T, Yuan J, Zaretsky JM, Desrichard A, Walsh LA, Postow MA, Wong P, Ho TS, et al. Genetic Basis for Clinical Response to CTLA-4 Blockade in Melanoma. N. Engl. J. Med. 2014;371(23):2189–2199. doi:10.1056/NEJMoa1406498

59. Kalaora S, Wolf Y, Feferman T, Barnea E, Greenstein E, Reshef D, Tirosh I, Reuben A, Patkar S, Levy R, et al. Combined Analysis of Antigen Presentation and T-cell Recognition Reveals Restricted Immune Responses in Melanoma. Cancer Discov. 2018;8(11):1366–1375. doi:10.1158/2159-8290.CD-17-1418

60. Gros A, Parkhurst MR, Tran E, Pasetto A, Robbins PF, Ilyas S, Prickett TD, Gartner JJ, Crystal JS, Roberts IM, et al. Prospective identification of neoantigen-specific lymphocytes in the peripheral blood of melanoma patients. Nat. Med. 2016;22(4):433–438. doi:10.1038/nm.4051

61. Prickett TD, Crystal JS, Cohen CJ, Pasetto A, Parkhurst MR, Gartner JJ, Yao X, Wang R, Gros A, Li YF, et al. Durable Complete Response from Metastatic Melanoma after Transfer of Autologous T Cells Recognizing 10 Mutated Tumor Antigens. Cancer Immunol. Res. 2016;4(8):669–678. doi:10.1158/2326-6066.CIR-15-0215

62. Parkhurst M, Gros A, Pasetto A, Prickett T, Crystal JS, Robbins P, Rosenberg SA. Isolation of T-Cell Receptors Specifically Reactive with Mutated Tumor-Associated Antigens from Tumor-Infiltrating Lymphocytes Based on CD137 Expression. Clin. Cancer Res. 2017;23(10):2491–2505. doi:10.1158/1078-0432.CCR-16-2680

63. Stevanović S, Pasetto A, Helman SR, Gartner JJ, Prickett TD, Howie B, Robins HS, Robbins PF, Klebanoff CA, Rosenberg SA, et al. Landscape of immunogenic tumor antigens in successful immunotherapy of virally induced epithelial cancer. Science. 2017;356(6334):200–205. doi:10.1126/science.aak9510

64. Le DT, Durham JN, Smith KN, Wang H, Bartlett BR, Aulakh LK, Lu S, Kemberling H, Wilt C, Luber BS, et al. Mismatch repair deficiency predicts response of solid tumors to PD-1 blockade. Science. 2017;357(6349):409–413. doi:10.1126/science.aan6733

65. Deniger DC, Pasetto A, Robbins PF, Gartner JJ, Prickett TD, Paria BC, Malekzadeh P, Jia L, Yossef R, Langhan MM, et al. T-cell Responses to TP53 “Hotspot” Mutations and Unique Neoantigens Expressed by Human Ovarian Cancers. Clin. Cancer Res. 2018;24(22):5562–5573. doi:10.1158/1078-0432.CCR-18-0573

66. The problem with neoantigen prediction. Nat. Biotechnol. 2017;35(2):97–97. doi:10.1038/nbt.3800

67. Chen DS, Mellman I. Elements of cancer immunity and the cancer–immune set point. Nature. 2017;541(7637):321–330. doi:10.1038/nature21349

68. Havel JJ, Chowell D, Chan TA. The evolving landscape of biomarkers for checkpoint inhibitor immunotherapy. Nat. Rev. Cancer. 2019;19(3):133–150. doi:10.1038/s41568-019-0116-x

69. Falk K, Rötzschke O, Stevanovié S, Jung G, Rammensee H-G. Allele-specific motifs revealed by sequencing of self-peptides eluted from MHC molecules. Nature. 1991;351(6324):290–296. doi:10.1038/351290a0

70. Abelin JG, Harjanto D, Malloy M, Suri P, Colson T, Goulding SP, Creech AL, Serrano LR, Nasir G, Nasrullah Y, et al. Defining HLA-II Ligand Processing and Binding Rules with Mass Spectrometry Enhances Cancer Epitope Prediction. Immunity. 2019;51(4):766-779.e17. doi:10.1016/j.immuni.2019.08.012

71. Davis MM. The problem of plain vanilla peptides. Nat. Immunol. 2003;4(7):649–650. doi:10.1038/ni0703-649

72. Song I, Gil A, Mishra R, Ghersi D, Selin LK, Stern LJ. Broad TCR repertoire and diverse structural solutions for recognition of an immunodominant CD8+ T cell epitope. Nat. Struct. Mol. Biol. 2017;24(4):395–406. doi:10.1038/nsmb.3383

73. Stewart-Jones >GBE, McMichael AJ, Bell JI, Stuart DI, Jones EY. A structural basis for immunodominant human T cell receptor recognition. Nat. Immunol. 2003;4(7):7.

74. Celie PHN, Toebes M, Rodenko B, Ovaa H, Perrakis A, Schumacher TNM. UV-Induced Ligand Exchange in MHC Class I Protein Crystals. J. Am. Chem. Soc. 2009;131(34):12298–12304. doi:10.1021/ja9037559

75. Turner SJ, Doherty PC, McCluskey J, Rossjohn J. Structural determinants of T-cell receptor bias in immunity. Nat. Rev. Immunol. 2006;6(12):883–894. doi:10.1038/nri1977

76. Wang GC, Dash P, McCullers JA, Doherty PC, Thomas PG. T Cell Receptor αβ Diversity Inversely Correlates with Pathogen-Specific Antibody Levels in Human Cytomegalovirus Infection. Sci. Transl. Med. 2012;4(128):128ra42. doi:10.1126/scitranslmed.3003647

77. Yager EJ, Ahmed M, Lanzer K, Randall TD, Woodland DL, Blackman MA. Age-associated decline in T cell repertoire diversity leads to holes in the repertoire and impaired immunity to influenza virus. J. Exp. Med. 2008;205(3):711–723. doi:10.1084/jem.20071140

78. Yarchoan M, Johnson BA, Lutz ER, Laheru DA, Jaffee EM. Targeting neoantigens to augment antitumour immunity. Nat. Rev. Cancer. 2017;17(4):209–222. doi:10.1038/nrc.2016.154

79. Łuksza M, Riaz N, Makarov V, Balachandran VP, Hellmann MD, Solovyov A, Rizvi NA, Merghoub T, Levine AJ, Chan TA, et al. A neoantigen fitness model predicts tumour response to checkpoint blockade immunotherapy. Nature. 2017;551(7681):517–520. doi:10.1038/nature24473

80. Wells DK, van Buuren MM, Dang KK, Hubbard-Lucey VM, Sheehan KCF, Campbell KM, Lamb A, Ward JP, Sidney J, Blazquez AB, et al. Key Parameters of Tumor Epitope Immunogenicity Revealed Through a Consortium Approach Improve Neoantigen Prediction. Cell. 2020;183(3):818-834.e13. doi:10.1016/j.cell.2020.09.015

81. June CH. Drugging the Undruggable Ras — Immunotherapy to the Rescue? N. Engl. J. Med. 2016;375(23):2286–2289. doi:10.1056/NEJMe1612215

82. Chen J-L, Stewart-Jones G, Bossi G, Lissin NM, Wooldridge L, Choi EML, Held G, Dunbar PR, Esnouf RM, Sami M, et al. Structural and kinetic basis for heightened immunogenicity of T cell vaccines. J. Exp. Med. 2005;201(8):1243–1255. doi:10.1084/jem.20042323

83. Miles JJ, Tan MP, Dolton G, Edwards ESJ, Galloway SAE, Laugel B, Clement M, Makinde J, Ladell K, Matthews KK, et al. Peptide mimic for influenza vaccination using nonnatural combinatorial chemistry. J. Clin. Invest. 2018;128(4):1569–1580. doi:10.1172/JCI91512

